# FtsK, a DNA motor protein, coordinates the genome segregation and early cell division processes in *Deinococcus radiodurans*

**DOI:** 10.1101/2022.07.15.500248

**Authors:** Shruti Mishra, Hari S. Misra, Swathi Kota

**Author notes:** Corresponding authors’ addresses Dr. Swathi Kota Molecular Biology Division, Bhabha Atomic Research Centre, Mumbai- 400094. India. Dr. Hari S. Misra Molecular Biology Division, Bhabha Atomic Research Centre, Mumbai-400094. India.

## Abstract

FtsK/SpoIIIE protein family are DNA translocases known as the fastest DNA motor proteins that use ATP for their movement on DNA. Most of the studies in single chromosome-containing bacteria have established the role of FtsK in chromosome dimer resolution (CDR) connecting the bacterial chromosome segregation process with cell division. But, only limited reports are available on the interdependent regulation of genome segregation and cell division in multipartite genome harbouring (MGH) bacteria. In this study, for the first time, we report the characterization of FtsK from the radioresistant MGH bacterium *Deinococcus radiodurans* R1 (drFtsK). drFtsK shows the activity characteristics of a typical FtsK/SpoIIIE/Tra family. It stimulates the site-specific recombination catalyzed by *Escherichia coli* tyrosine recombinases. drFtsK interacts with various cell division and genome segregation proteins of *D. radiodurans*. Microscopic examination of different domain deletion mutants of this protein reveals alterations in cellular membrane architecture and nucleoid morphology. *In-vivo* localization studies of drFtsK-RFP show that it forms multiple foci on nucleoid as well as on the membrane with maximum density on the septum. drFtsK coordinates its movement with nucleoid separation. The alignment of its foci shifts from old to new septum indicating its cellular dynamics with the FtsZ ring during the cell division process. Nearly, similar positional dynamicity of FtsK was observed in cells recovering from gamma radiation exposure. These results suggest that FtsK forms a part of chromosome segregation, cell envelope, and cell division machinery in *D. radiodurans*.

**Importance:** *Deinococcus radiodurans* show extraordinary resistance to γ-radiation. It is polyploid and harbours a multipartite genome comprised of two chromosomes and two plasmids, packaged in a doughnut-shaped toroidal nucleoid. Very little is known about how the tightly packed genome is accurately segregated and the next divisional plane is determined. FtsK, a multifunctional protein, helps in pumping the septum-trapped DNA in several bacteria. Here, we characterized FtsK of *D. radiodurans* R1 (drFtsK) for the first time and showed it to be an active protein. The absence of drFtsK causes many defects in morphology at both cellular and nucleoid levels. The compact packaging of the deinococcal genome and cell membrane formation is hindered in *ftsK* mutants. *In-vivo* drFtsK is dynamic, forms foci on both nucleoid and septum, and coordinates with FtsZ for the next cell division. Thus, drFtsK role in maintaining the normal genome phenotype and cell division in *Deinococcus radiodurans* is suggested.

## 1. Introduction

Bacteria orchestrate their cell division in time and space by the coordinated action of proteins involved in cell division named divisome and genome maintenance named segrosome. The divisome assemblies contain more than 20 proteins in a variety of bacteria like *Escherichia coli*, *Neisseria gonorrhoeae, Pseudomonas aeruginosa, etc* (Aarsman et al., 2005; Goehring and Beckwith, 2005; Pazos et al., 2013) which ensures an accurate cell constriction, septal peptidoglycan (PG) synthesis, and finally cell separation (den Blaauwen et al., 2017). One of these proteins, FtsK was first documented in *Escherichia coli* TOE44 conditional mutant that was impaired in cell division. It was observed that at 42°C, its mutant forms highly indented filaments suggesting blockage at a very late stage of division and hence was named <u>F</u>ilament temperature-<u>s</u>ensitive mutant K. TOE44 (AB2497 ftsK44) (Begg et al., 1995). Later, cellular localization studies revealed FtsK foci formation on the septum of *E. coli* (Yu et al., 1998) and this localization of FtsK along with FtsA recruits other cell division proteins like FtsL, FtsQ and FtsI, thus is required for the stabilization of divisome formed at the mid-cell position (Goehring et al., 2005). FtsK was characterized as a multifunctional DNA translocase belonging to the AAA+ (ATPase Associated with various cellular Activities) superfamily proteins (Aussel et al., 2002). Not only in the stabilization of divisional septa and DNA translocation but it is also involved in membrane synthesis and cell envelope remodeling by interacting with various proteins like septal peptidoglycan binding protein rare lipoprotein A (RlpA) and proteins involved in peptidoglycan synthesis like FtsI (Berezuk et al., 2018; Di Lallo et al., 2003). The protein is divided into three domains-the N-terminal domain of around 200 amino acids (NTD), the linker region (FtsKL), and the 500 amino acids C-terminal domain (CTD) (Draper et al., 1998; Grainge, 2010). In *E. coli*, the NTD is responsible for the attachment to the cell membrane by transmembrane (TM) segments and the interaction with other cell division and cell envelope proteins (Berezuk et al., 2014). The FtsKL is not conserved and varies in length and composition between species (Bigot et al., 2007, Dubarry et al., 2010). The CTD is highly conserved and comprises three separate subdomains called α, β, and γ (Yates et al., 2003). This region is known for the motor functions of FtsK and helps in DNA translocation. The γ domain (FtsKγ) is attached to the β-domain via a flexible linker and recognizes 8-bp sequences called KOPS (FtsK Orienting Polar Sequence, GGGNAGGG) in *E. coli* and SRS (SpoIIIE recognition sequence, GAGAAGGG) in *B. subtilis*. KOPS/SRS sequences are asymmetrically distributed on the chromosome arms and they direct the translocation of FtsK/SpoIIIE/Tra family proteins towards the “*dif*” (deletion-induced filamentation) site in the “*ter”* region (Levy et al., 2005; Becker et al., 2007). In this region, two ‘*dif*’ sites are brought closer by FtsK which thereby recruits and activates tyrosine recombinases XerCD to form the synaptic complex XerCD-*dif* (Ip et al., 2003). This XerCD-*dif* complex is involved in site-specific recombination (SSR) to resolve the chromosome dimers (chromosome dimer resolution-CDR) (Grainge et al., 2011; Keller et al., 2016). In this way, FtsK helps in sorting duplicated chromosome copies at the site of cell division (Lesterlin et al, 2005).

*Deinococcus radiodurans* R1 is a coccus-shaped gram-positive bacterium that shows extreme resistance to multiple abiotic stresses like gamma radiation, desiccation, and oxidative stress (Cox and Battista, 2005; Makarova et al., 2001). These phenotypes are attributed to its highly efficient DNA double-strand break repair and a strong anti-oxidant mechanism (Slade and Radman, 2011; Misra et al., 2013). This bacterium exists in tetrads and consists of a multipartite genome system (MGS) with two chromosomes (Chr I, Chr II) and two plasmids [Mega plasmid (MP) and small plasmid (SP)]. It has a polyploid multipartite genome, which is highly catenated and intertwined making the nucleoid tightly arranged in the form of a very distinct doughnut-shaped toroidal structure (Levin-Zaidman et al., 2003). How this genome arrangement is maintained and segregated during cell division is still unknown. The *D. radiodurans* genome encodes putative FtsK/SpoIIIE family protein (White et al., 1999). In what way FtsK might act to help segregate multiple, complex chromosomes is an outstanding question of great interest. Here, we report the characterization of FtsK of *D. radiodurans* (drFtsK) and its potential role in genome separation, septum formation, and cell division. We demonstrated that the purified drFtsK showed concentration-dependent ATPase activity, sequence-specific interaction with KOPS DNA of *E. coli,* and could stimulate SSR by *E. coli* XerCD *in-vitro*. Protein-protein interaction studies indicated that drFtsK interacts with various genome segregation and cell division proteins. Further, different domain deletion mutants of dr*ftsK* exhibited a significant change in the growth rate under normal conditions as well as after irradiation during the post-irradiation recovery (PIR) period. Morphological studies indicated that the deletion of different domains of drFtsK affected the nucleoid arrangement and cellular architecture in this bacterium resulting in some abnormal phenotypes. The localization studies of FtsK-RFP expressed under native promoter showed the appearance of dispersed fluorescent foci on the nucleoid, septal and peripheral membranes of the cells and a coordinated movement with FtsZ during cell growth. Together, these results suggest that drFtsK helps in cross-talks between cellular and molecular events like genome segregation, cell envelope remodeling, and septum formation in *D. radiodurans*.

## 2. Materials and methods

### 2.1 Bacterial strains, growth conditions, plasmids, and materials

All the bacterial strains and plasmids used in this study are listed in Table S1. *Deinococcus radiodurans* R1 (ATCC13939) was a kind gift from Professor J. Ortner, Germany (Schaefer et al., 2000). This bacterium was grown in TYG (Tryptone (0.5%), Yeast extract (0.3%) and Glucose (0.1%) medium at 32°C. *Escherichia coli* strains were grown in Luria-Bertini (LB) broth (1% tryptone, 0.5% yeast extract, and 1% sodium chloride) at 37°C. *Escherichia coli* strain DH5α was used for cloning purposes and maintenance of all the plasmids. For the expression of recombinant proteins, *E. coli* strain BL21 (DE3) pLysS was used. Cells harbouring pET28a (+) derivatives or Bacterial Two-Hybrid System (BACTH) vectors and their derivatives were maintained in the presence of the respective antibiotic selection pressure. Shuttle expression vector pVHSM (Charaka and Misra, 2012) and its derivatives were maintained in the presence of spectinomycin in *E. coli* and *D. radiodurans* cells (70 μg/ml and 40 μg/ml, respectively). Deinococcal cells harbouring suicide vector-pNOKOUT (Khairnar NP et al., 2008) derivatives were grown in the presence of kanamycin (8μg/ml). All the enzymes and molecular biology grade chemicals were purchased from Sigma Chemical Company, USA, and Merck India Pvt. Ltd. India. Isopropyl-β-D-1-thiogalactopyranoside (IPTG), DAPI, Nile red, and vancomycin hydrochloride (unlabelled) were purchased from Merck Inc. Vancomycin BODIPY FL conjugate (labeled) was purchased from Invitrogen. Syto-green9 dye was purchased from Sigma Aldrich. Antibody against T18 (SC-33620) domain of CyaA of *Bordetella pertussis*, was obtained commercially from Santa Cruz Biotechnology, Inc. Antibody against polyhistidine tag was procured from Sigma. Radiolabelled nucleotides were acquired from the Board of Radiation and Isotope Technology, Department of Atomic Energy (DAE), India (BRIT, India).

### 2.2 Bioinformatic analysis and molecular modeling

The *D. radiodurans* genome encodes a putative FtsK (locus tag-E5E91_ RS02025, old locus tags-Dr_0400 and Dr_0401) on chromosome I, annotated as DNA translocase FtsK (hereafter called drFtsK). The full-length FtsK sequence of *D. radiodurans* and its other characterized homologs in *B. subtilis* (SpoIIIE_Bs), *E. coli* (FtsK_Ec), and *P. aeruginosa* (FtsK_Pa), *S. aureus* (FtsK_Sa), *V. cholerae* (FtsK_Vc) were retrieved from the NCBI genome database. Multiple sequence alignment and secondary structure prediction were performed with the Promals3D online server (Pei et al., 2008). The neighbor-joining phylogenetic tree (without distance corrections) between deinococcal FtsK and known FtsK family proteins was constructed using the PHYLIP program. After alignment, the Walker box (walker A ‘P loop’ and walker B) ATP-binding motif and DNA-binding motif were searched. Structural models of drFtsK protein domains (motor domain and gamma domain) were constructed by the I-TASSER server (http://zhanglab.ccmb.med.umich.edu /I-TASSER/). The models were validated by the Swiss model workspace. Templates used for the modelling of drFtsK domain structures were derived from the known structures of respective *E. coli* FtsK motor domain (PDB ID: 2ius) and gamma domain (PDB ID: 2j5p). The modeled structures of the deinococcal FtsK domains were superimposed with *E. coli* FtsK domain structures (2ius and 2j5p) using Pymol software and RMSD and TM-score were analyzed which indicated high similarity between the models. KOPS motifs were searched across the *D. radiodurans* genome by DistAMo online tool, shown in Figure S4 (Sobetzko et al., 2016).

### 2.3 Cloning, expression, and purification of recombinant proteins

A list of all the primers used for constructing recombinant plasmids and generating deletion/insertion mutants are mentioned in Table S2. The open reading frames (ORF) of drFtsK (Dr_0400/0401), drFtsKΔN (Dr_0400), drFtsKγ, ecFtsKγ, ecXerC, ecXerD were PCR amplified from *D. radiodurans* or *E. coli* genome (as applicable) using sequence-specific primers. PCR products were purified and ligated at *Nde*I and *BamH*I sites in pET28a (+) to produce the respective plasmids (Table S1). These plasmids were transformed into *E. coli* BL21 (DE3) pLysS. Induction of expression and preparation of cell-free extract was performed as reported earlier (Kota and Misra, 2006). Purification of proteins was done by nickel affinity chromatography. For that, the cell-free extract was loaded onto NiCl_2_ charged-fast-flow-chelating-sepharose column (GE Healthcare) which was pre-equilibrated with buffer (20mM Tris-HCl, 300mM NaCl, 10% glycerol) containing 10 mM imidazole. Extensive washing of column was done with 20 volumes of buffer A having 50 mM imidazole. Step-wise elution of recombinant proteins was done using 100 mM, 200 mM, and 250 mM imidazole in buffer A and analysis of each fraction was carried out on 10% SDS-PAGE. Fractions containing the desired proteins at the expected size were pooled and dialyzed in buffer A containing 100 mM NaCl and proceeded for HiTrap™ Heparin HP column purification. drFtsKΔN protein quality was examined by Circular dichroism (CD) spectroscopy in buffer (20 mM Tris-HCl, pH 7.6, 100 mM NaCl) using JASCO, J815 with ∼0.2 mg/ml, as described earlier (Modi et al. 2014). The oligomeric status of the protein (5μM) was checked by Dynamic Light Scattering (DLS) in the same buffer as mentioned before using the Malvern Panalytical Zetasizer nano range instrument with/without KOPS DNA (100nM) and ATP (0.5mM).

### 2.4 ATPase activity assay

ATPase activity detection was done by thin-layer chromatography method to track the release of [^32^P]-αADPs from [^32^P]-αATPs and by malachite green assay to quantify the amount of inorganic phosphate (Pi) released as described in (Modi et al. 2014). Briefly, different concentrations of drFtsKΔN were added with 30 nM [^32^P] -α ATP in a buffer containing 20 mM Tris (pH7.6), 75 mM KCl, 2 mM MgCl_2_ and incubated at 37°C for 0.5 hours The reaction was discontinued using 10 mM EDTA and 1µl of each reaction mixture was spotted on the PEI-Cellulose F+ TLC sheet. Components were separated on solid support after being air-dried in a buffer containing 0.75M KH2PO4/H3PO4 (pH 3.5), and an autoradiogram was developed. For the colorimetric quantitative assay, ATPase/GTPase activity assay kit (sigma-Aldrich) was used with/without KOPS DNA (100nM), and µmole Pi was released per min per µl was calculated. Data were plotted using the GraphPad Prism 6 software.

### 2.5 Protein-DNA interaction study

The DNA-binding activity of drFtsKΔN was checked by electrophoretic mobility shift assay (EMSA) as described in (Leonard, 2005). In brief, ssDNA containing *E. coli dif* and KOPS (72mer) or *E. coli dif* (40mer) sequence were radiolabelled with [^32^P]-γATP using T4 polynucleotide kinase (as mentioned in Löwe et al., 2008). Forward strands were annealed with the complementary strands to yield radiolabeled dsDNA (*dif* and *dif*+KOPS). Approximately 30nM radiolabelled dsDNA was incubated with different concentrations of drFtsKΔN (0–2µM) in a buffer containing 50 mM Tris-Cl (pH 8.0), 75 mM NaCl, 5 mM MgSO4 and 0.1 mM DTT for 30 min at 37°C with/without 2mM ATP. The reaction mixture was loaded on 8% native PAGE gel, the gel was dried and the autoradiogram was developed.

### 2.6 Recombination assay

An *in-vitro* recombination reaction was performed as reported by (Löwe et al., 2008). 1 nM radiolabelled *dif* fragment and 10 nM dif-KOPS fragment was incubated with 150 nM ecXerC, 30 nM ecXerD in buffer containing 10 mM MgCl_2,_ 20 mM Tris/HCl pH 7.6. To this, 40 nM FtsK variants (drFtsKΔN, drFtsKγ, and ecFtsKγ) were added separately and incubated for 2 min. The reaction was monitored in the presence/absence of 2 mM ATP for 30 min at 37°C. The reaction was stopped by adding 0.1% SDS and 0.1 mg/ml proteinase K and the mixture was run through 7% denaturing PAGE gel containing 0.1% SDS in 1X TBE buffer. The gel was dried and an autoradiogram was developed. Band intensities on the autoradiogram were estimated densitometrically by ImageJ 2.0 software and % recombination product was calculated.

### 2.7 Generation of drftsK deletion mutants and ftsK-rfp knock-in mutant

For creating deletion mutants of the different domains of FtsK in *D. radiodurans*, the respective regions of the coding sequence of *ftsK* (Figure S6) were replaced by *nptII* as described earlier (Khairnar et al., 2008). In brief, ∼1 kb upstream and ∼1 kb downstream sequences to the target region were PCR amplified from *D. radiodurans* genomic DNA using sequence-specific primers (as given in Table S2). The upstream fragment was cloned at *Apa*I and *EcoR*I sites while the downstream fragment was cloned at *BamH*I and *Xba*I sites into a suicide pNOKOUT vector (Khairnar et al., 2008) to produce pNKFKUD, pNKFK_MC_UD, pNKFK_N_UD plasmids (Table S1). The recombinant plasmids were linearized with *Xmn*I and transformed into *D. radiodurans* separately. Transformants obtained were plated on TGY plates containing kanamycin (8µg/ml) and grown for many generations under necessary antibiotic pressure to obtain homozygous insertion of *nptII* and replacement of the target portions of *ftsK* in the whole deinococcal genome. This was confirmed by PCR amplification using *ftsK* domain-specific primers as well as antibiotic (*npt*II) cassette-specific primers (Table S2). The homozygous replacement of the target genes with the *nptII* cassette was accomplished and these cells with genotype *ftsK*::*nptII, ftsKN*::*nptII, ftsKMC*::*nptII* were denoted as Δ*ftsK,* Δ*ftsKN,* Δ*ftsKMC* respectively. The *ftsKN*::*nptII* mutant has a deletion of N-terminal 1-163 amino acids of drFtsK (M1-L163), *ftsKMC*::*nptII* mutant has a deletion of 295-1209 amino acids of drFtsK (D295-K1209) and *ftsK*::*nptII* has a deletion of full-length drFtsK (M1-K1209).

For expression of *ftsK-rfp* under the native promoter of *ftsK*, the replacement of chromosomal copy of *ftsK* with *ftsK-rfp* was carried out using a similar approach as mentioned above. The translational fusion of FtsKγ-RFP was generated by cloning the *ftsK*-gamma domain (*ftsK*γ) in pDsRed vector to yield pDsRedFKγ. Further, *ftsKγ-rfp* (from pDsRedFKγ) was cloned upstream to *nptII* and downstream *ftsk* sequence was cloned downstream to *nptII* in pNOKOUT to yield pNKFKγRD. This was transformed into *D. radiodurans* to obtain cells with genotype *ftsK::ftsK-rfp* which expressed FtsK-RFP under the native promoter.

### 2.8 Growth studies of different deletion mutants of ftsK

*D. radiodurans* R1 wild-type and *ftsK* mutants were exposed to 6kGy gamma-radiation as described in (Misra et al., 2006). Briefly, overnight grown bacterial cultures (with/without kanamycin; 8 µg/ml) were washed and suspended in sterile phosphate-buffered saline (PBS). These cells were then exposed to 6 kGy γ-radiation at a dose rate of 1.5 kGy/h (Gamma Cell 5000, ^60^Co, Board of Radiation and Isotopes Technology, DAE, India). Equal numbers of irradiated (IRR) and control un-irradiated cells (UI) were grown in TYG medium (with/without kanamycin; 8 µg/ml) in 96 well microtiter plates after washing with PBS (Nunclon; Sigma-Aldrich). Optical density at 600nm was measured to examine the growth in replicates at 32 °C for 18 h using Synergy H1 Hybrid multi-mode microplate reader. The growth curves were fitted using the spline regression model. Statistical analysis was carried out using the statistical programs R, http://www.r-project.org/, the “spline” package was used to generate the spline regression-based models, in R. The model was developed using linear b-splines with two knots producing three intervals-1,2,3. Fixed time intervals were chosen as 0-5hrs (1), 5-10hrs (2), and 10-18hrs (3). The linear growth rates were computed by the change in absorbance (OD) per interval. Un-irradiated samples were compared with R1-UI as control and irradiated samples were compared with R1-IRR as control. The generation time of wildtype and *ftsk* mutants were calculated as mentioned in (Maier, 2009). Graphs of growth rate coefficient v/s strain at each time interval and generation time were plotted using Graphpad Prism 6.0. Significance value (P-value) obtained at 95% confidence intervals were considered to be significantly different.

### 2.9 Microscopy studies of different deletion mutants of ftsK

Confocal microscopy was done on IX3SVR using an Olympus IX83 inverted microscope with the laser beams focused on the back focal plane of a 100 × 1.40 NA oil-immersion apochromatic objective lens (Olympus) as described earlier (Maurya et al., 2019). The time sequence and intensity of laser illumination at samples were adjusted using the installed FLUOVIEW software. For imaging, a series of Z-planes were acquired at every 400-nm using a motorized stage, and then z-stacking was done to create 3D images. In brief, *D. radiodurans* R1 wild-type cells and different deletion mutants of *ftsK* in exponential phase at three-time points-6 hrs, 9 hrs, and 14 hrs were taken (Figure 5 and S7). The cells were washed twice with phosphate-buffered saline (pH 7.4). These cells were stained with Nile red (1 mg/ml) for membrane and DAPI (40,6-diamidino-2-phenylindole, dihydrochloride) (0.2 mg/ml) for genome for 10 min on ice and then washed thrice with PBS. Cells were then mounted on a 1% agarose bed on glass slides and imaged. Fluorescence from Nile Red/ RFP tagged protein was detected using TRITC (tetramethylrhodamine isothiocyanate) filter with 542/562nm, excitation/emission wavelength. Fluorescence from DAPI was detected using its filter with 402/460nm, excitation/emission wavelength. Each image is represented in separated as well as merged channels. The brightness and contrast of all images were tuned using Adobe Photoshop 7.0. For quantification of image patterns, 200-300 cells from two-three isolated microscopic fields were taken in independent experiments and analyzed for necessary attributes. Image analysis and other cell parameters were determined using automated Olympus CellSens software. Data obtained was plotted using Graphpad Prism 6.0.

### 2.10 Localization of FtsK and FtsZ in wild-type and ΔftsK mutant

For studying the localization of FtsK-RFP foci, unlabelled vancomycin and labeled (Van-FL-BIODPY) were combined in the ratio of 1:1 (0.5 µg/ml) to stain the membrane as mentioned before (Tiyanont et al., 2006) and exponential phase cells expressing FtsK-RFP under native promoter were grown for 90-120 min. These cells were washed twice with phosphate-buffered saline (pH 7.4) and stained with DAPI. Fluorescence from Van-FL-BODIPY/ GFP tagged protein was detected using a FITC filter with 475/530, excitation/emission wavelength. Image analysis was done as described above. Around 150 diad and 150 tetrad cells were counted separately for the quantification of localization of FtsK-RFP foci on the nucleoid / peripheral membrane or the outer membrane. Data were plotted as a scatter plot in GraphPad Prism 6 software. For FtsZ localization studies, the recombinant plasmid pVHZGFP expressing FtsZ-GFP was transformed in wild-type (WT) and Δ*ftsK*-mutant cells of *D. radioduran.* Cells were grown overnight and cells equivalent to 0.05-0.1 OD_600_ were diluted with a fresh medium containing the required antibiotic and induced with 10mM IPTG overnight.

### 2.11 Growth phase-dependent studies on FtsK and FtsZ dynamics

Time-lapse imaging was done for the monitoring of the dynamics of FtsK with respect to genome movement, cell growth, and division. Cells expressing FtsK-RFP were stained with 150 nM of Syto-green9 dye and PBS washed with cells were placed on an agarose pad made in 2X-TYG and constructed with air holes for oxygenation of cells. Fluorescence emission was grabbed with DM-488/561 dichroic mirror and corresponding single-band emission filters at different time intervals. Images were taken for a period of 4 hr at intervals of 1 hour using very low laser power (561 nm and 488 nm). For monitoring FtsK localization and dynamics under gamma radiation exposure, the cells were treated with 6kGy radiation and grown for 1hr with constant shaking. Then, the cells were visualized as mentioned above. We did a line scan analysis (LSA) for each time point through CellSens software. In LSA, we scanned the fluorescence intensity of Syto-green and FtsK-RFP signals across the growing/developing septum to find the relative intensity of the genome and FtsK-RFP. For studying FtsK and FtsZ dynamics, cells expressing FtsK-RFP under native promoter were transformed with pVHZGFP and similarly proceeded for time-lapse microscopy as explained above. All the co-localization studies were done using built-in CellSens Software and the represented co-localization patterns showed Pearson’s correlation coefficient (R(r)) >0.7-1.0 and Overlap coefficient (R) of 0.9-1.0. Co-localization was represented as white foci in the images. 76 cells were analyzed at each time point to determine the number of FtsK foci co-occurring with FtsZ. Values obtained were plotted using Graphpad prism-6.0.

### 2.12 Protein-protein interaction study

The possible interaction of drFtsK with other deinococcal proteins of cell division like FtsZ, FtsA, and DivIVA, and genome segregation proteins like TopoIB, ParB2, ParB3, and ParB4 was monitored in surrogate *E. coli* by using the co-immunoprecipitation method. The recombinant pUT18 plasmids were already used in the earlier studies (Maurya et al., 2016, Maurya et al., 2018, Chaudhary et al., 2019, Kota et al., 2021) were transformed in *E. coli* BL21 (DE3) pLysS strain expressing Histidine tagged FtsK (pETFtsK). *E. coli* cells co-expressing histidine-tagged FtsK with T18-tagged protein in separate combinations were obtained at the log phase and cell-free extracts were prepared. Total proteins were immunoprecipitated using anti-polyhistidine antibodies and the possibility of T18 tagged partners’ presence in immunoprecipitate was checked using monoclonal antibodies against the T18 domain of CyaA as depicted earlier (Maurya et al., 2016, 2018). Hybridization signals were detected with anti-mouse secondary antibody labeled with alkaline phosphatase using NBT/BCIP substrates (Roche Biochemical, Mannheim).

### 2.13 Statistical analysis

All the statistical analysis was done using-Student’s t-test. Significance value (P value) obtained at 95% confidence intervals are depicted as **** for P value <0.0001, *** for P value <0.001, ** for P value of 0.05-0.001, * for P value <0.05.

## 3. Results

### 3.1 The *D. radiodurans* genome encodes an FtsK homologue

Upon BLAST search, we found that the putative deinococcal FtsK (drFtsK) was wrongly annotated in the genome of *D. radiodurans*. Indeed, the coding sequence of putative drFtsK spans upon two Open Reading Frames (ORF) i.e. Dr_0400 and Dr_0401 in chromosome I. Dr_0401 contains the N-terminal region (229 amino acids) and Dr_0400 contains the remaining region (980 aa) (White et al., 1999). Recently, this has been corrected and the entire coding sequence of FtsK is annotated with a locus tag- E5E91_ RS02025 (Repar et al. 2021). Multiple sequence alignment using the PROMALS3D tool showed that drFtsK has ∼25-30% identity with FtsK/SpoIIIE members of other species [*E. coli* (FtsK_Ec), *V. cholerae* (FtsK_Vc), *P. aeruginosa* (FtsK_Pa), *L. lactis* (FtsK_Ll), *S. aureus* (FtsK_Sa), *B. subtilis* (SpoIIIE_Bs)]. The overall protein homology is less because the N-terminal and linker region is quite variable but the C-terminal region is highly conserved (Figure 1A and Figure S1A). Consensus sequences for ATP binding P-loop motif (Walker A motif), Walker B motif and winged-winged helix-turn-helix (wHTH) DNA binding region were compared with the drFtsK sequence. The results showed that the C-terminal domain of drFtsK contains a consensus ATP binding motif- “GSTGSGKS” and DNA binding motif - “HARAGKLMDLLR” indicating it is an FtsK homologue (Figure 1A). Further, the phylogenetic analysis revealed that drFtsK forms a separate clade (Figure S1B). Domain prediction exhibited that drFtsK contains the canonical FtsK domains- FTSK_4TM (FtsKN), FtsK_alpha, FtsK_SpoIIIE (αβ motor pump) and FtsK_gamma domain (drFtsKγ) (Figure 1B). Modeled structures of the drFtsK gamma and motor domains were aligned with the template structures of gamma (2j5p) and motor (2ius) domains in *E. coli* FtsK, respectively. The corresponding aligned models showed RMSD values of 0.21 (TM score 0.938) and 0.34 (TM score 0.962), respectively, indicating that these structures are highly identical (Figure 1C). This information led us to hypothesize that drFtsK may perform similar functions as that of characterized FtsK/SpoIIIE proteins from other bacteria provided that this bacterium has the functional homologs of tyrosine recombinases. We also searched for putative tyrosine recombinases taking *E. coli* XerCD-specific domains for analysis. Six putative ORFs, in chromosome I- E5E91_RS02620 (old locus tag- Dr_0513); in chromosome II- E5E91_RS13705 (old locus tag- Dr_A0075), E5E91_RS14115 (old locus tag- Dr_A0155) and E5E91_RS14250 (old locus tag- Dr_A0182); in megaplasmid- E5E91_RS15640 (old locus tag- Dr_B0104) and in small plasmid- E5E91_RS15930 (old locus tag- Dr_C0018) were identified (Figure S2). Which combination of these ORFs would function as tyrosine recombinases and is significant in genome segregation in this bacterium is worth understanding and will be addressed separately.

**Figure 1:**
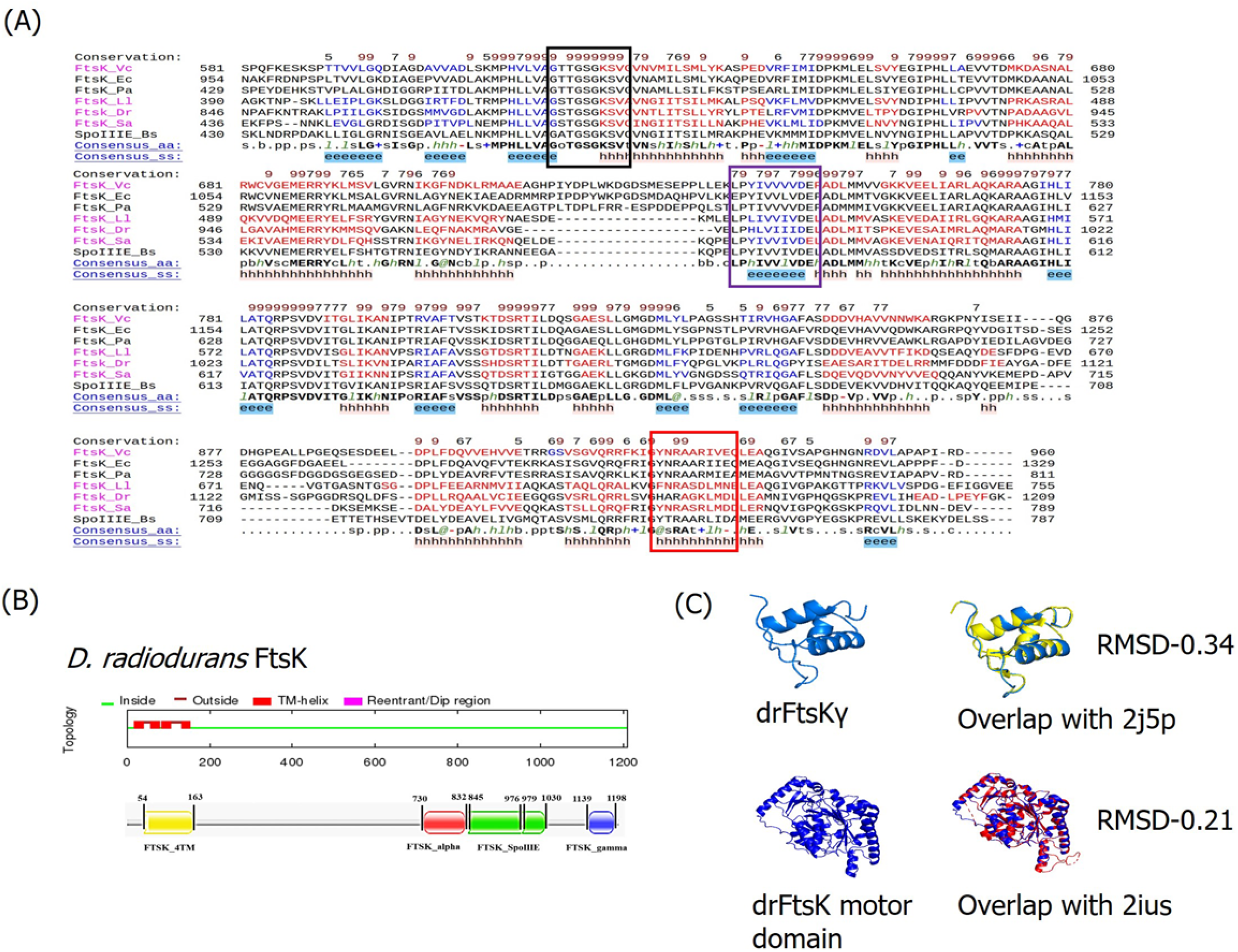
Deinococcal FtsK protein sequence alignment and modeling. (A) Multiple sequence alignment (MSA) of drFtsK with known FtsK/SpoIIIE family proteins. The amino acid sequences of FtsK from *Deinococcus* r*adiodurans* (FtsK_Dr), SpoIIIE from *Bacillus subtilis* (SpoIIIE_Bs), FtsK of *Escherichia coli* (FtsK_Ec), *Pseudomonas aeruginosa* (FtsK_Pa)*, Lactococcus lactis* (FtsK_Ll)*, Vibrio cholerae* (FtsK_Vc), and *Staphylococcus aureus* (FtsK_Sa) are collected from NCBI and the homology between sequences is checked by PROMALS3D webserver. MSA of the C-terminal region is depicted here. Boundaries of the conserved motifs in the C-terminal are marked as a black box for the walker A domain (ATP binding P-loop motif), a purple box for the Walker B motif, and a red box for the DNA binding motif. Predicted secondary structures are displayed below the sequences. (B) Different domains present in drFtsK are represented as FTSK_4TM (FtsK_N terminal region), FtsK_alpha, FtsL_SpoIIIE (αβ motor pump) and FtsK_gamma domain(drFtsKγ). (C) Modeled structures of drFtsKγ and motor domain were aligned with the template structures of *E. coli* FtsK-gamma domain (2j5p) and *E. coli* FtsK motor domain (2ius), respectively.

### 3.2 drFtsK shows ATPase activity

The N-terminal truncated drFtsK contains all functional domains like ATP and DNA interacting motifs, and gamma domain and lacks transmembrane segments that would be required for localization and *in-vivo* function. So, N-terminal truncated drFtsK (drFtsKΔN) could be used for biochemical characterization *in-vitro*. Therefore, we checked the ATPase activity and DNA binding activity of purified drFtsKΔN. Circular dichroism (CD) analysis showed that purified drFtsKΔN does contain folded protein and the protein has majorly α-helical conformation (Figure 2A), which was supported by *in-silico* secondary structure prediction by PROMALS3D (Figure S1A). Further, the oligomeric status of the protein was checked by dynamic light scattering. Results showed that the protein exits in hexameric form even in absence of DNA (Figure S3A). This indicates that drFtsK binds to DNA in a pre-formed hexameric form as shown previously for SpoIIIE in *B. subtilis* (Cattoni et al., 2013). The ATPase activity of purified drFtsKΔN was determined using [^32^P]-αATP hydrolysis (Figure 2B) as well as a colorimetric malachite green ATPase activity assay (Figure 2C). Results showed that drFtsKΔN is an ATPase that could convert ATP to ADP and inorganic phosphate. An increase in the release of inorganic phosphate was seen with the increase in protein concentration. However, the specific activity of the enzyme remained unaltered. Further, the presence of *E. coli* KOPS containing DNA could marginally stimulate the ATPase activity of purified drFtsKΔN (Figure S3B).

**Figure 2:**
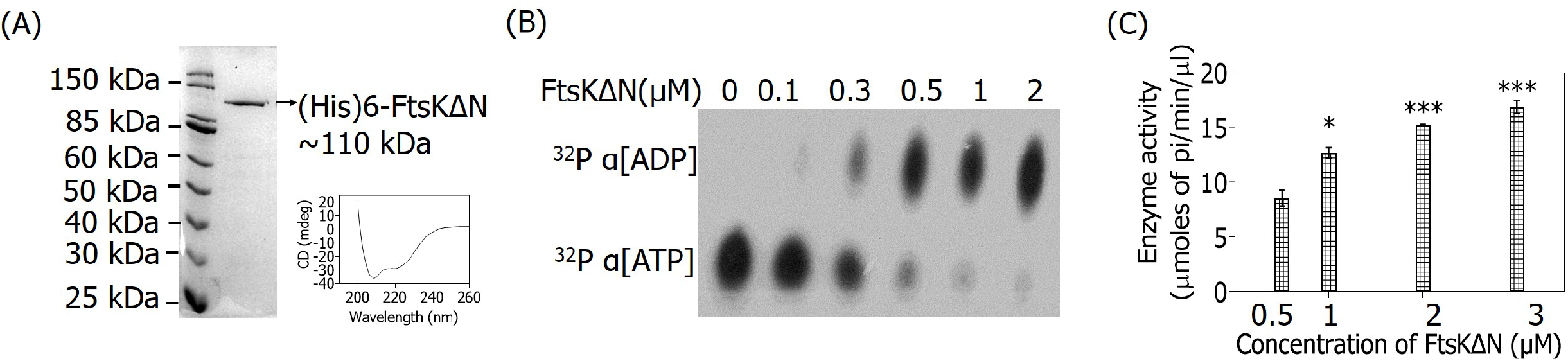
The ATPase activity of drFtsK. (A) Polyhistidine tagged fusion protein of N-terminal truncated protein (FtsKΔN) was purified and the purified protein was checked for proper refolding by circular dichroism as described in the methods. (B) ATPase activity of recombinant purified FtsKΔN protein was checked using radiolabelled ATP [^32^P]-αATP. Different concentrations of the protein were incubated with radiolabelled nucleotide and reaction products were separated on PEI-Cellulose F^+^ TLC. The autoradiogram shows the hydrolysis of ^32^P-α[ATP] to ^32^P-α[ADP]. (C) Quantitative analysis of ATPase activity was done by colorimetric malachite green reagent using increasing protein concentration. Data shown here are the mean+-SD (n=3) plotted in GraphPad Prizm6. Statistical significance was obtained by student’s t-test. The p-values attained at 95% confidence intervals are depicted as (***) for <0.001 and (*) for <0.05.

### 3.3 drFtsK binds to Escherichia coli KOPS and activates site-specific recombination catalyzed by tyrosine recombinases -XerCD

It has been known that the FtsK protein of *E. coli* (ecFtsK) binds to KOPS and the orientation of these motifs on the chromosome is highly skewed so that they direct the FtsK translocase towards the terminal replichore region for decatenation of the duplicated circular chromosome. Several key residues implicated in KOPS recognition have been identified in the ecFtsK gamma domain (N1296, R1300, E1303; Sivanathan et al., 2006). We found that these residues are not identical across FtsK/SpoIIIE family proteins, which could lead to differences in the FtsK recognition sites in different organisms. Therefore, we analyzed the distribution of a few KOPS octamers in the deinococcal genome using DistAMo online tool. The results indicated that there is a high frequency of GGGNAGGG motif family and *E. coli* KOPS-GGGCAGGG on deinococcal chromosomes I with high density existing near the ‘ter’ region. Compared to chromosome I, chromosome II has less density (Figure S4). Thus, over- or under-representation of these motifs suggests that the GGGCAGGG motif might function as KOPS in *D. radiodurans.* But, compared to *E. coli,* the overall frequency of the GGGCAGGG motif is very less in the *D. radiodurans* genome. However, we monitored the DNA binding activity of drFtsKΔN by EMSA using radiolabelled dsDNA sequences containing *E. coli dif* + KOPS. The protein showed a sequence-specific binding with dsDNA containing the GGGCAGGG motif (Figure 3B). The size of the nucleoprotein complex increased gradually (reflected as slower mobility) with the rise in protein concentration suggesting more FtsK molecules are binding to KOPS. Earlier, it was shown that in general, three FtsKγ domains bind to 8 bp KOPS DNA (Löwe et al., 2008). The presence of ATP did not affect drFtsKΔN binding with KOPS sequences. This shift is not observed in the case of dsDNA containing only the *E. coli dif* site (Figure 3A) suggesting that drFtsK interaction is specific to GGGCAGGG-KOPS that could act as the loading site for drFtsK on the genome in *D. radiodurans*.

**Figure 3:**
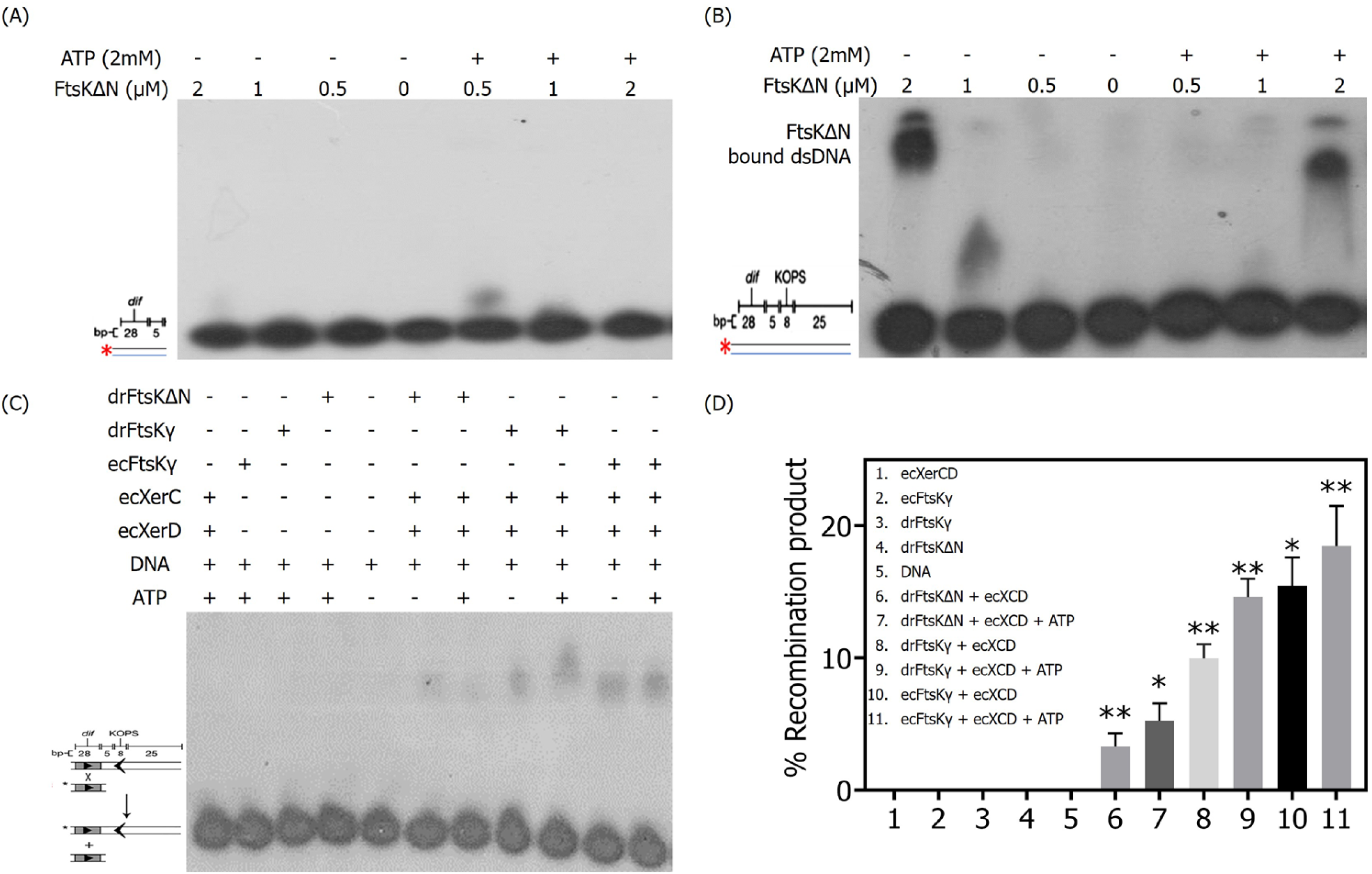
DNA binding activity and activation of *E. coli* tyrosine recombinases by deinococcal FtsKΔN protein. Purified recombinant FtsKΔN protein was checked for DNA binding activity with [*γ*-32P] ATP labelled dsDNA containing *E. coli dif*-40bp (A), *E. coli dif* and KOPS-72bp (B) in the presence/absence of ATP. Autoradiograms show EMSA gels where no interaction of drFtsKΔN is seen with only *E. coli dif* sequence but interaction with *E. coli dif* and KOPS is seen. A schematic representation of the recombination reaction substrate (short radiolabelled *dif* containing dsDNA sequence) and product (long *dif* + KOPS containing dsDNA sequence) as described earlier is given (Löwe et al., 2008) (C). Autoradiogram showing site-specific recombination reaction by *E. coli* tyrosine recombinases XerC and XerD (ecXerCD). Recombination products were obtained in those reactions where FtsK (drFtsKΔN/ drFtsKγ/ ecFtsKγ) was present with EcXerCD. Band intensities obtained by densitometric analysis of the autoradiogram were used to calculate % recombination product in each reaction (D). Data shown here are the mean+-SD (n=3) and statistical significance was found using the student’s t-test. The p-values attained at 95% confidence intervals are depicted as (*) for <0.05 and (**) for 0.05-0.001.

As drFtsKΔN showed binding with KOPS, we tested the ability of drFtsKΔN and drFtsKγ to activate XerCD site-specific recombination (SSR). For this, the well-characterized *E. coli* recombination system components were utilized and an *in-vitro* recombination assay was carried out using purified ecXerC (34kDa) and ecXerD (36kDa) proteins (Figure S5) as described previously (Löwe et al., 2008) and represented in Figure 3C. Results showed that drFtsKΔN and drFtsKγ can stimulate ecXerCD mediated recombination between two dsDNA substrates-one containing only *dif* sites and the other containing *dif +*KOPS (Figure 3C). Both drFtsKΔN and drFtsKγ showed a nearly similar pattern of recombination products in the presence and absence of ATP. The recombination at *dif* sites in the presence of drFtsKΔN and drFtsKγ proteins could be possible only when these proteins get loaded at KOPS and then translocate through dsDNA until it reaches the *dif* site and stimulates recombination activity of ecXerCD there. Upon quantification of the percent recombination product, it was noticed that the recombination reaction has occurred at low efficiency in the presence of drFtsKΔN. With drFtsKγ, it was found to be nearly similar to that observed with ecFtsKγ (recombinant protein containing *E. coli* FtsK gamma domain) which was used as a positive control (Figures 3D and 3E). These results together suggested that drFtsKΔN and drFtsKγ proteins are functional at least *in-vitro* and exhibit ATPase, *E. coli* KOPS binding, and stimulate tyrosine recombination by *E. coli* XerCD at *E. coli dif* sites.

### 3.4 FtsK deletion affects growth rate and morphology in D. radiodurans

To understand the *in-vivo* roles of drFtsK, we created chromosomal deletions of the different regions in the coding sequences of this protein, and the effects of these deletions on growth and morphology were examined. The *D. radiodurans* cells containing the deletion of full-length FtsK (Δ*ftsK*), middle and C-terminal domain (*ΔftsKMC),* and N-terminal domain (*ΔftsKN)* (as depicted in Figure S6) showed a differential effect on growth under normal un-irradiated conditions (Figure 4). For instance, Δ*ftsKN* behaved nearly similar to wild type (Figure 4A) while *ΔftsKMC* and *ΔftsK* mutants showed extended lag period (Figure 4B and 4C). These mutants also showed a significant decrease in the growth rate under un-irradiated conditions and after gamma irradiation (6kGy) treatment (Figure 4D). The effect of drFtsK domains deletion on the growth rate is growth phase-dependent. The growth rate of *ftsK* mutants-*ΔftsKMC* and *ΔftsK* are slower than the wildtype (R1) and Δ*ftsKN* initially (time interval 0-5hrs) but is faster at the later stages (time interval 10-18hrs). The decreased rate was observed particularly during the lag and exponential phases where cellular processes like genome duplication, segregation, and cell division are actively occurring. This resulted in a decrease in the generation time of the mutants under un-irradiated and after gamma irradiation treatment also (Figure 4E). The slow growth rate in mutants indicated that FtsK plays some critical role in normal cellular processes like genome segregation and cell division in this bacterium.

**Figure 4:**
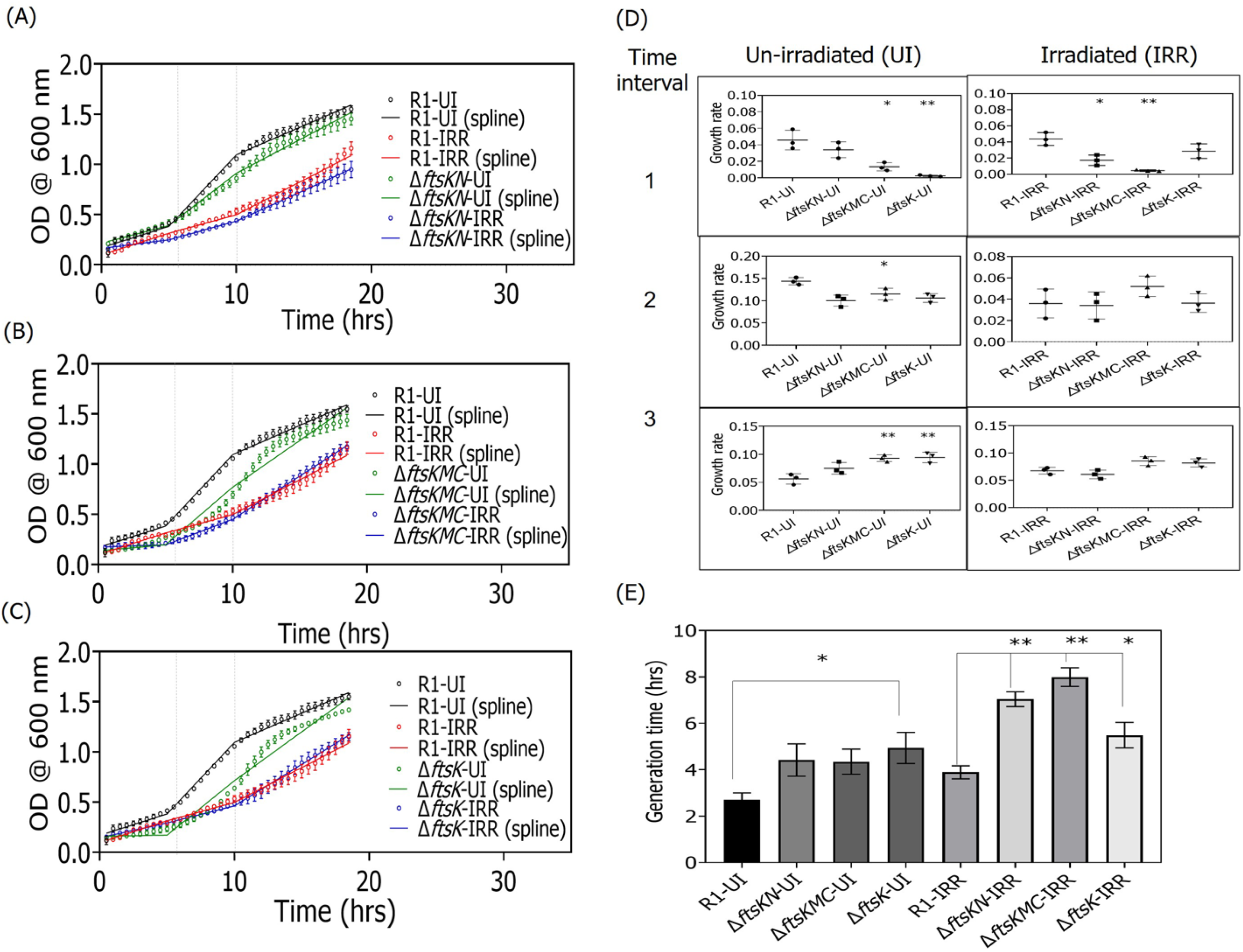
Effect of deinococcal FtsK deletion on the growth of *D. radiodurans.* Cell survival studies of different deletion mutants of *ftsk -ΔftsKN* (A)*, ΔftsKMC* (B)*, and ΔftsK* (C) were monitored under un-irradiated conditions (normal) and after gamma irradiation treatment (6kGy). The growth data (circles) were fitted by a linear spline regression model (lines). The dashed vertical lines show the knots dividing the spline modeled growth curve into three intervals. The difference in the growth rates of wild-type (R1) and different domain deletion mutants under normal and post-irradiation recovery conditions at each time interval was calculated by estimating the slopes of each line segment from the spline regression (D). The generation time (doubling time) of wildtype and *ftsk* mutants under normal and irradiated conditions were calculated as described in the methodology and plotted (E). Data shown here are the mean+-SD (n=9) and the statistical significance of the differences was found using the student’s t-test. The p-values attained at 95% confidence intervals are depicted as (*) for <0.05 and (**) for 0.05-0.001.

Further, we compared the cell morphology of *ftsK* deletion mutants with wild-type (WT) under non-irradiated (normal) conditions at 14 hrs post subculturing. In *ΔftsKN*, *ΔftsKMC,* and *ΔftsK* mutants, although a majority of the cells showed typical tetrad and nucleoid arrangement, around ∼10% cell population showed a significant change in their phenotypes as compared to wild-type (Figure 5A and 5B). Mainly, three types of atypical phenotypic changes were observed in these mutants; abnormal tetrad arrangements (ABN), bent septum (BS), and anucleated cells within the tetrad (AC-T). Besides these, other parameters like cell diameter, nucleoid compaction (as determined by nucleoid diameter), nucleoid intensity and percent of tetrads and diads in a population were also observed to be statistically different in all three mutants (Figure 5C, 5D, 5E, and 5F). For example, the nucleoid diameter is more in mutants than in wild type. Similar observations were found with the other phenotypes scored. To understand it better, we expressed FtsZ-GFP (pVHZGFP) in *ΔftsK* mutant and observed that a significant population (∼10%) in *ftsK* deletion mutant got affected in the temporal and spatial regulation of FtsZ functions in *D. radiodurans* as seen by FtsZ-GFP ring misplacement (Figure 5G). Earlier, it was shown that deletion of the C-terminal region of DivIVA protein in *D. radiodurans* generates the bent septum (Chaudhary et al., 2021). So, the presence of bent septum in *ΔftsK* mutant may be attributed to its probable role in the positioning of the pole determining protein and divisome components in this bacterium. Not only bent septum and defective FtsZ ring formation but the FtsK mutants formed membrane bulges and showed changes in membrane staining properties at exponential phase time points-6 & 9 hrs of subculturing indicating the role in membrane biology in *D. radiodurans* (Figure S7). The *ΔftsKN* mutants were able to mostly recover from this at 9 hrs but the other two mutants could not. Further, the survival of cells upon *ftsK* deletion even after significant phenotypic changes might argue its functional redundancy in this bacterium.

**Figure 5:**
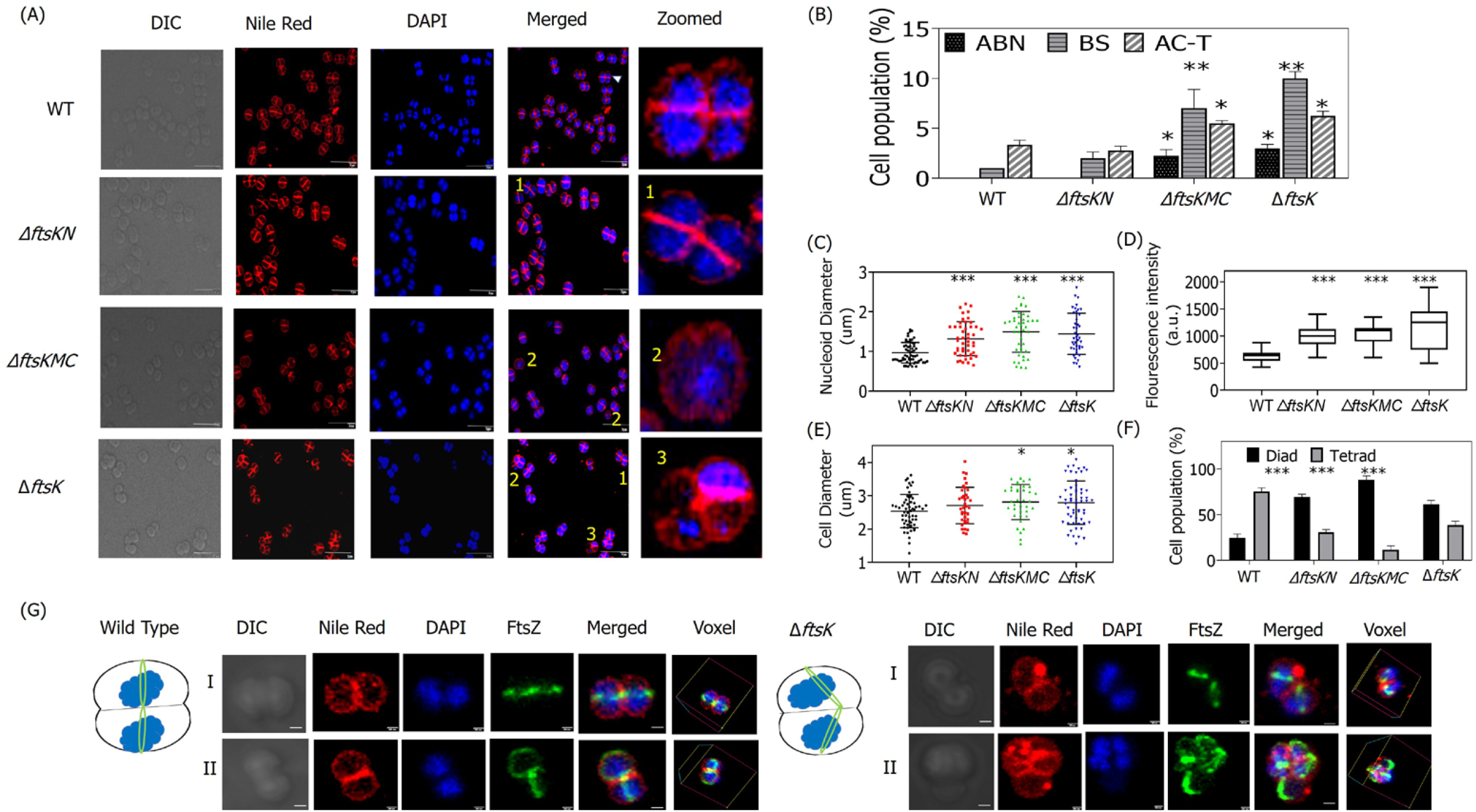
Effect of different domain deletion of *drftsK* on the phenotype of *D. radiodurans* cells. Cells at 14 hrs post subculturing was taken and fluorescence microscopic images show the DIC, TRITC (for Nile Red), DAPI, and merged channels of the wild type (WT), and different mutants of *ftsk*-*ΔftsKMC*, *ΔftsKN, ΔftsK* cells (scale bar-5µm). A significant population of cells in *ΔftsKMC*, *ΔftsK, and ΔftsKN* mutants showed a change in tetrad cell morphology, nucleoid arrangement, and septal membrane. Atypical phenotypes are represented as; 1. Bent septum (BS), 2. Abnormal tetrad arrangements (ABN), and 3. Anucleated cells within the tetrad (AC-T) were obtained (zoomed cell scale bar-500nm) (A). Statistical analysis of the different morphologies obtained in *ftsK* mutants as compared to WT cells is done in 200-300 cells and plotted (B). Other phenotypic changes like nucleoid diameter (C), nucleoid DAPI fluorescence intensity (D), cell diameter (E), and percent of diads or tetrads in the cell populations (F) were calculated and plotted. The p-values attained at 95% confidence intervals are depicted as (*) for <0.05, (**) for 0.05-0.001, (***) for <0.001. Microscopic images of the cell division septal ring formation by drFtsZ-GFP in WT and *ΔftsK* mutant cells are depicted along with pictorial representation (scale bar-500nm) (G). The images shown here are representative pictures of the experiments conducted at least three times.

### 3.5 FtsK localizes on nucleoid and septum in D. radiodurans

The translational fusion of FtsK-RFP was expressed under a native promoter in *D. radiodurans* and the cellular localization of drFtsK was observed microscopically. Interestingly, FtsK-RFP produced foci on the membrane (septal and peripheral) and the nucleoid (Figure 6A). When we quantitated the number of FtsK-RFP foci in the cell, the majority of the foci were found to be on the septal membrane (SM) in comparison with the peripheral membrane (PM) and nucleoid (N), both in the diad and tetrad population (Figure 6B). This result became more apparent when bacterial growth was relatively slowed down by growing it on the agar plate and observing it under the microscope. Maximum FtsK-RFP foci density was seen on the septum (Figure 6C). Around 200 cells were analyzed to determine the fraction of cells showing localization at the old or new septum, dispersed; and both on the new and old septum making a cross pattern (Figure 6D). The formation of the new septum was determined based on the vancomycin staining or the appearance of septum constriction as observed from the DIC pictures. Based on these observations, it can be speculated that FtsK protein localization is dynamic in *D. radiodurans* showing on both nucleoid and septum depending on the growth stage of the cells.

**Figure 6:**
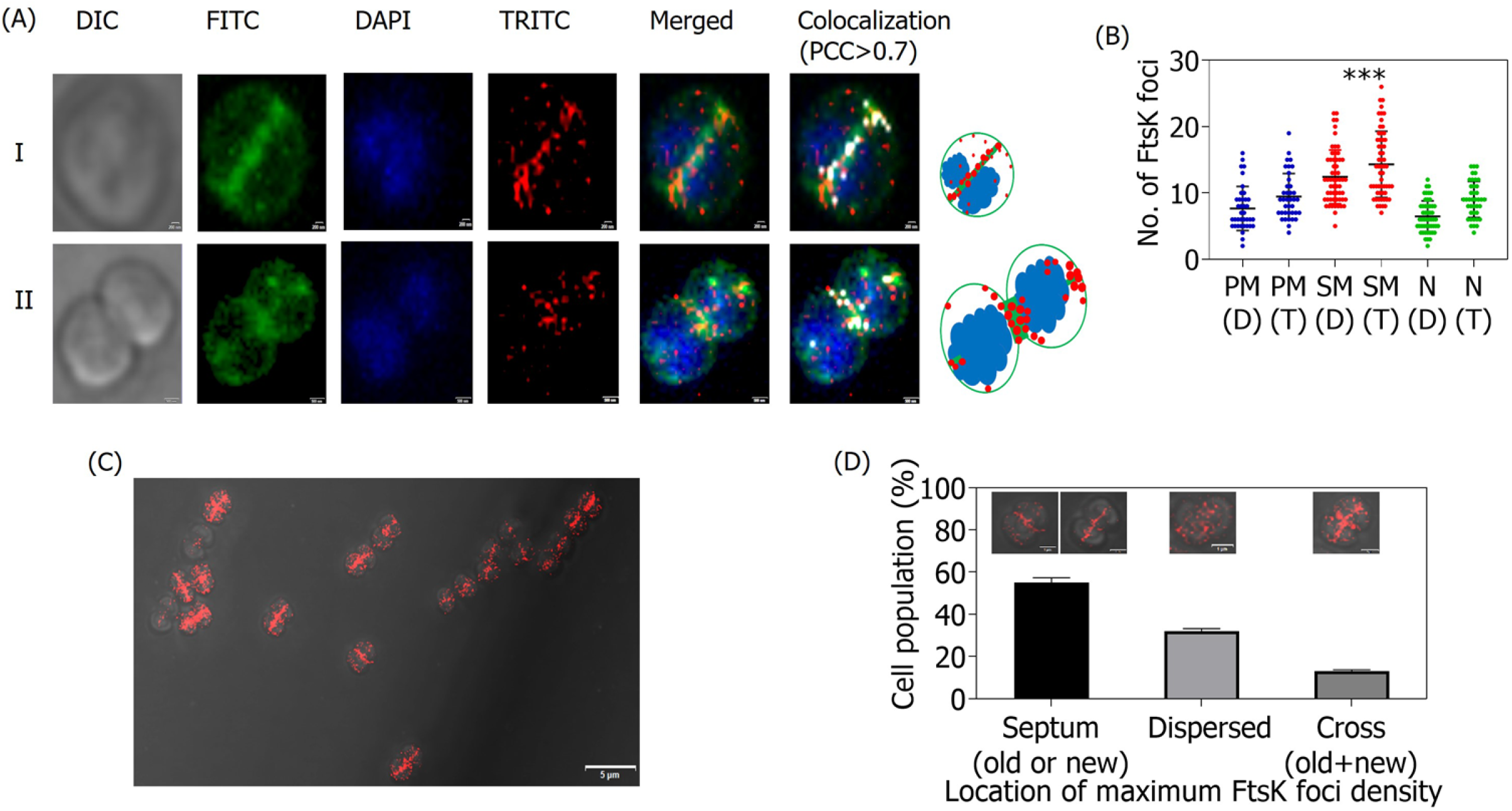
Cellular localization of FtsK-RFP expressing under native promoter in *D. radiodurans cells*. Expression of FtsK-RFP foci (TRITC channel-red) in the cells is seen after nucleoid staining by DAPI (blue) and membrane staining by Vancomycin-BioDIPY (FITC channel-green). FtsK-RFP shows association on the nucleoid as well as on the membrane, both septal and peripheral membrane. Data represented here shows microscopic images of two independent cells along with pictorial images (scale bar-200nm and 500nm for I and II, respectively) (A). White foci show the co-localization of FtsK-RFP with the membrane (PCC>0.7). Diad and tetrad cells were counted separately for the location of FtsK-RFP foci on the peripheral membrane (PM-D for diad and PM-T for tetrad), on the septal membrane (SM-D for diad and SM-T for tetrad), and on nucleoid (N-D for diad and N-T for tetrad. A significantly high foci density is seen on the septum as analyzed in both diads and tetrads (B). The p-values attained at 95% confidence intervals are depicted as (***) for <0.001. Cells grown in stationary conditions show FtsK-RFP expression in different locations-foci on the old or new septum, dispersed foci, and foci on both old and new septum making a cross pattern (scale bar-5μm) (C). The percentage of cells showing different cellular localization of FtsK-RFP foci in a population of ∼200 cells was calculated and plotted (D). The images shown here are representative pictures of the experiments conducted at least three times.

### 3.6 Time-lapse microscopy shows the movement of FtsK-RFP along the new septum in D. radiodurans

We observed that FtsK-RFP localization on the nucleoid and the septum is heterogenous possibly because of the non-synchronous population of the cells. This prompted us to do time-lapse microscopy to observe the FtsK dynamics in dividing cells of *D. radiodurans*. Time-lapse microscopy was performed under both normal growth and the post-irradiation recovery phase (Figure 7). Under normal conditions, at time point t=0, maximum FtsK-RFP foci are aligned at the septum. With progression in the cell division; the alignment of the foci shifts to the newly forming septum perpendicular to the old septum (t=2). This alignment is seen when the nucleoid has separated into two and the septum constriction has just initiated. By t=4, FtsK-RFP foci are seen to be moving to align along the next probable plane of division (Figure 7A). In gamma-radiation treated cells, the transition in the alignment of FtsK foci from the old septum to the new septum as the nucleoid divides is more apparent (Figure 7B). At t=0, FtsK-RFP foci are arranged along the membrane mostly with prominently high foci density at the initiation sites of the new pole on septal and peripheral membranes. As time proceeds, the majority of FtsK-RFP foci position along the new division septum perpendicular to the old one. Notably, line scan analysis (LSA) showed that the progression of FtsK alignment with the nucleoid separation process is very well synchronized (Figure 7B). Thus, FtsK-RFP dynamically switches from old septa to new septa as the cell division progresses under both normal and irradiated conditions. Orientation of FtsK at the septum would be functionally essential to activate CDR so that the genome can be separated properly and segregated correctly into daughter cells. It was observed that the nucleoid divides into two daughter toroidal structures simultaneously with the FtsK dynamics from the old septum to the new septum. *In E. coli*, FtsK is known to bind to KOPS sequences on the genome and translocates towards the *dif* site. It also acts as a translocase by pumping out the septum-trapped DNA during chromosome separation and cell division (Grainge, 2010). Our results obtained from the localization studies of drFtsK-RFP suggest a similar type of function in *D. radiodurans* also. Interestingly, there is an evident increase in the level of FtsK protein expression after irradiation. This could be explained by the transcriptome sequencing analysis where an increase in the *ftsK* transcript levels was seen after irradiation (Mishra et al., 2019). *D. radiodurans* contains 6-8 copies of the genome which are damaged when treated to 6kGy gamma radiation. So, the participation of translocases like FtsK during the repair and resolution of the multipartite genome cannot be ruled out. The increase in chromosome dimers possibly during extensive synthesis of DNA during post-irradiation recovery may require FtsK for activation of CDR in the cells. These results suggested that FtsK is dynamic and its dynamicity is required for the smooth and coordinated linking of genome segregation with early cell division processes like divisional plane determination and septum formation in this bacterium both under normal and post-irradiation recovery growth conditions.

**Figure 7:**
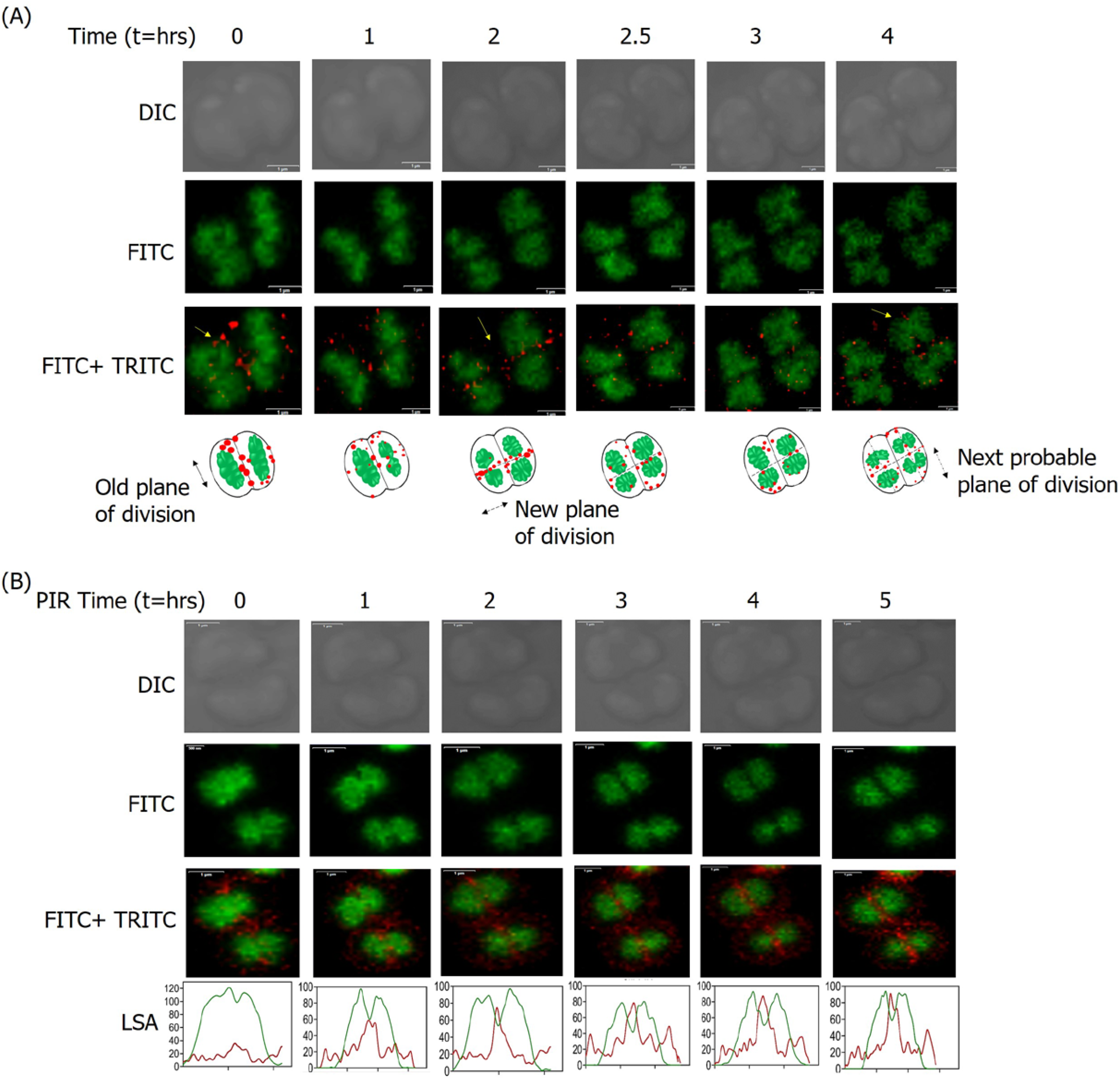
Cellular dynamics of deinococcal FtsK-RFP in normal and post-irradiation conditions. Time-lapse confocal microscopic images show FtsK-RFP (red foci) expressing cells stained with syto9 green dye (green) during the normal condition (A) and during post-irradiation recovery (PIR) (B). The top panels indicate the cell in planar view at different time points (t=hr) in the DIC channel, the middle panels indicate the genome arrangement in the FITC channel and the bottom panels show the same cell with TRITC and FITC channel merged to see FtsK-RFP foci dynamics at different ‘t’ (scale bar-1µm). The dynamics of FtsK-RFP foci is schematically represented under normal condition. During PIR, line scan analysis (LSA) shows the increase in the intensity of FtsK-RFP along the emerging septum.

### 3.7 Co-ordinated dynamics of FtsZ and FtsK in D. radiodurans

The septum localization of FtsK-RFP and the presence of bent septum in the deletion mutant of *ftsK* led us to study the dynamics of FtsK with another cell division protein-FtsZ concurrently in *D. radiodurans.* For this, the cells expressing FtsK-RFP under the native promoter were expressed with FtsZ-GFP on the pVHZGFP plasmid. During time-lapse microscopy, ∼75 cells were monitored for studying FtsZ and FtsK dynamics and analyzed. A single cell is shown in Figure 8A. At the start of imaging (t=0), the FtsZ ring is not complete but as the cell growth proceeds, the FtsZ ring is formed in ∼3 hours (t=3). The majority of FtsK-RFP can be seen aligned around the division septa at t=0.5 and then moving to position itself along the growing FtsZ ring at t=3 (Figure 8B). This suggests that drFtsK localizes to the newly forming divisional septum after the FtsZ ring formation is completed. These microscopic results showed that FtsK aligns along the developing septum acting in coordination with FtsZ. Further, co-localization analysis of FtsZ and FtsK proteins suggested that both the proteins significantly co-localize in some places. Our microscopic results are the first reports showing coordinated dynamics of the important cell division proteins i.e. FtsZ and FtsK in *D. radiodurans*.

**Figure 8:**
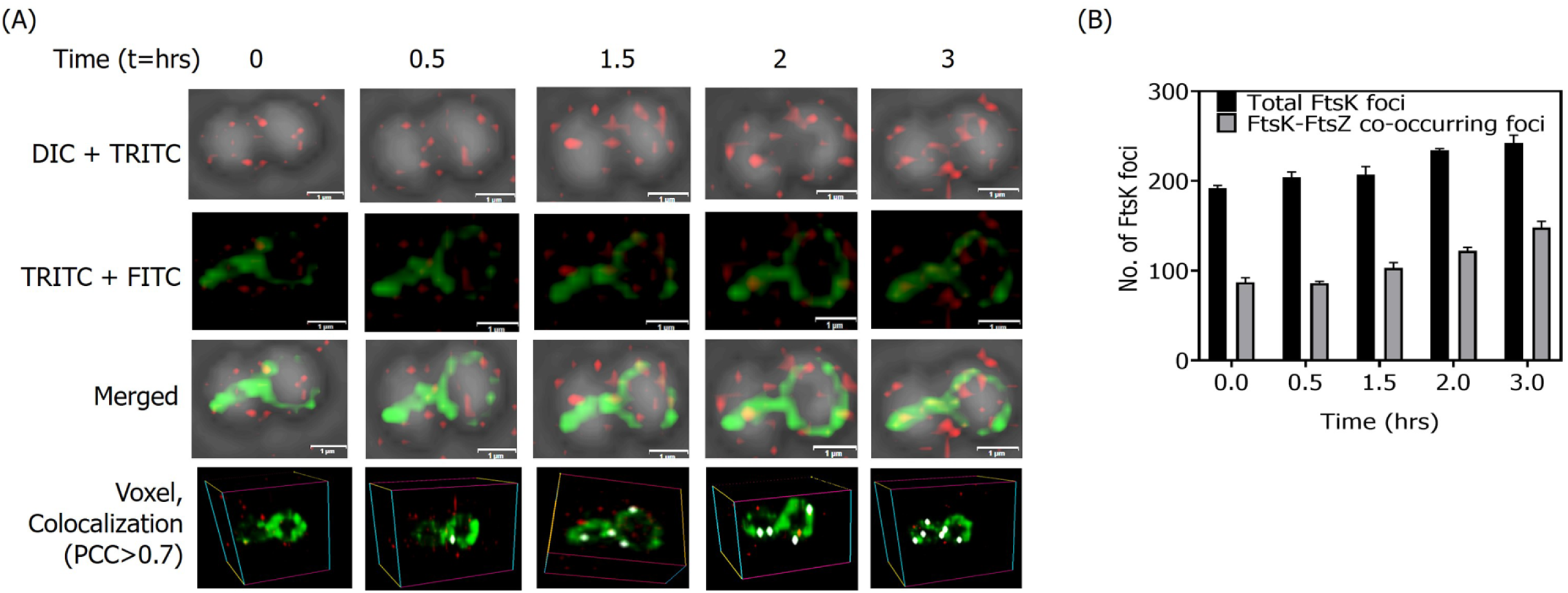
Co-ordinated cellular dynamics of cell division proteins FtsK and FtsZ in *D. radiodurans*. Time-lapse fluorescence microscopic images of cells expressing both FtsK-RFP and FtsZ-GFP show the dynamics of both the proteins in dividing cells under normal conditions. The top panel indicates the cell in planar view at different time points (t=hr) in the DIC+TRITC, the middle panels indicate the FITC+TRITC channel and merged images, and the bottom-most panel shows the same cell in voxel view with white foci showing the co-localization of FtsZ and FtsK (PCC>0.7) (scale bar-1µm). The majority of FtsK-RFP (red) can be seen aligned at the division septa at t=0.5hr and then moving to position themselves along the almost complete FtsZ ring at t=3hr (A). The total number of FtsK foci and FtsK-FtsZ co-occurring foci calculated in a population of ∼75 cells at time points of 0, 0.5h, 1.5h, 2h, and 3h were calculated and plotted (B).

### 3.8 FtsK interacts with divisome and segrosome components of D. radiodurans

Results obtained so far suggest that FtsK is dynamic during different stages of bacterial growth and plays an important role in both genome segregation and early cell division. The molecular basis of its dynamicity mainly relating to cell division and genome segregation would be worth understanding. Further, FtsK in other bacteria is known to interact with segrosome and divisome proteins. Therefore, the interaction of drFtsK with cell division and chromosome segregation proteins of *D. radiodurans* was checked by co-immunoprecipitation. It was observed that drFtsK interacts with deinococcal genome segregation proteins-ParB2, ParB3, ParB4, and TopoIB; cell division proteins -FtsZ, FtsA, and pole determining protein-DivIVA (Figure 9). Its interaction with DivIVA and FtsZ could explain the reason behind the bent septum phenotype observed in *ftsK* mutants. Surprisingly, it does not interact with some of the deinococcal proteins like ParAs, GyrA, FtsW, and FtsE which are involved in nucleoid segregation and cell division. These results suggested that drFtsK along with other genome segregation and cell division proteins may be involved in macromolecular complex formation for its overall functions and the possibility of this driving the dynamicity of FtsK independently of FtsZ cannot be ruled out in *D. radiodurans*.

**Figure 9:**
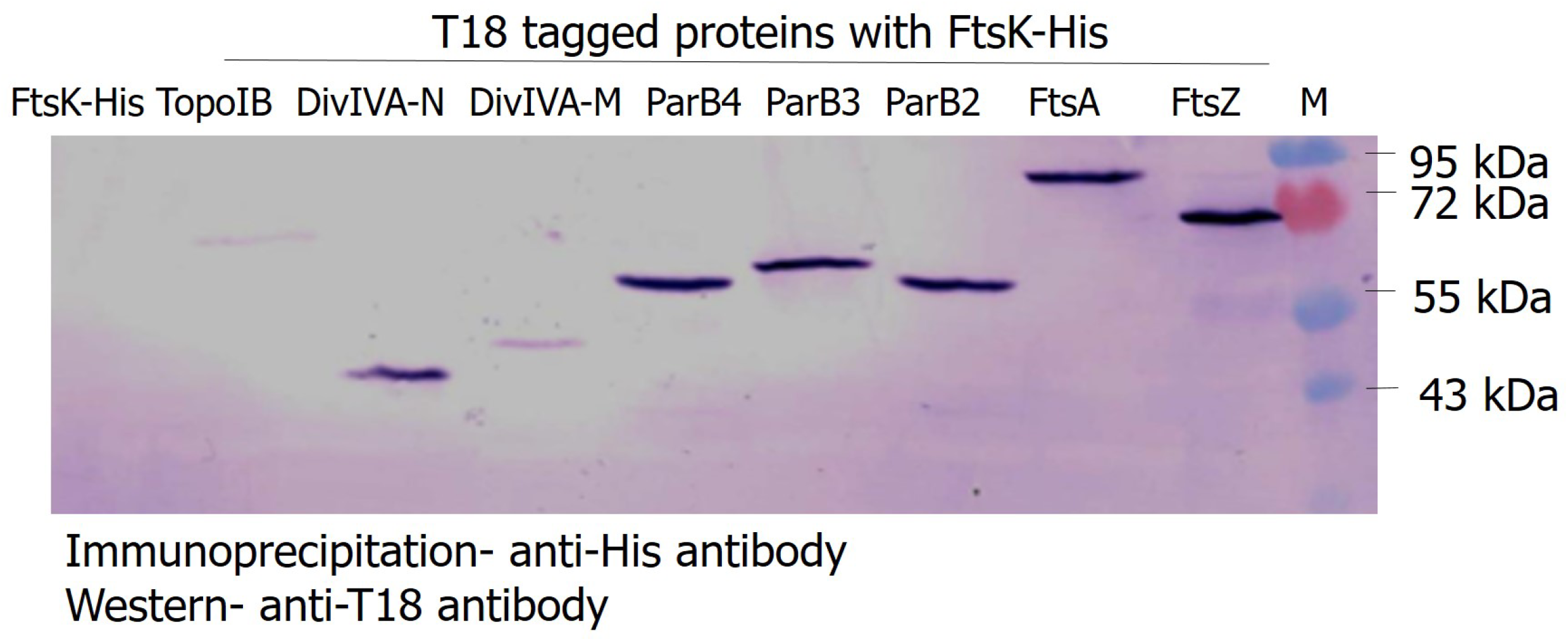
Interaction studies of deinococcal FtsK. Interaction of drFtsK and other proteins of *D. radiodurans* by co-immunoprecipitation assay followed by western blot. His-tagged drFtsK and T18-tagged ParB2, ParB3, ParB4, TopoIB, FtsA, DivIVA, and FtsZ in *E. coli* strain BL21 were checked for protein-protein interactions by co-immunoprecipitation using antibodies against polyhistidine-tag followed by immunoblotting using antibodies against T18 domain of CyaA. His-tagged FtsK expressing cells harbouring only the pUT18 vector was taken as a negative control. The experiment was conducted three times and the representative picture was shown.

## 4. Discussion

*Deinococcus radiodurans* R1 is a multipartite genome harbouring (MGH) bacterium having two chromosomes and two plasmids of 6-8 copies each per cell. The genome exists in a highly compacted nucleoid form which has been recently shown to dynamically change its shape during the cell division cycle (Floc’h et al. 2019). *D. radiodurans* exists in tetrads and grows by alternate planes of division where new cell division occurs in a plane perpendicular to the previous one. FtsK/SpoIIIE DNA translocases in bacteria are known to be involved in chromosome dimer resolution and pumping the trapped DNA through the newly formed septum during the cell division or sporulation. Despite the critical role of FtsK, it is reported to be dispensable for division under certain conditions (Goehring et al, 2007, Geissler and Margolin, 2005). Among MGH bacteria, the studies on FtsK have been reported in *V. cholerae* (Val et al., 2008; Midonet et al., 2014). Due to the complexities of multiple genome components in MGH, the role of FtsK in them would be fascinating to study. Our genome analysis revealed that *D. radiodurans* encodes putative FtsK. Unlike *V. cholerae*, where both the chromosomes exist separately in the cells, the entire genetic material in *D. radiodurans* is packaged in the form of a doughnut-shaped toroidal nucleoid. Therefore, the possibility of FtsK functioning differently in maintaining the genome integrity of *D. radiodurans* cannot be ruled out. Here, we have brought forth some proof to indicate that drFtsK is active and is involved in both genome segregation and early cell division processes (Figure 10). We demonstrated that recombinant FtsK is an ATPase with *E. coli* KOPS binding activity and could activate *E. coli* tyrosine recombinases XerCD *in-vitro* suggesting that drFtsK is functional in stimulating SSR. Fluorescence microscopy of cells expressing drFtsK-RFP showed the formation of multiple foci on nucleoid as well as on membrane. Multiple foci on nucleoid could be attributed to the presence of KOPS on the genome while foci formation on the membrane and /or septum could be attributed to FtsK transmembrane domain interacting with divisome components. A similar distribution of FtsK foci (∼30 to 100 molecules of FtsK per cell) was reported in *E.coli* (Bisicchia et al., 2013). Further, the localization pattern of FtsK in *D. radiodurans* seems to be cell cycle-specific where maximum protein intensity was observed on the septum in the dividing cells.

**Figure 10:**
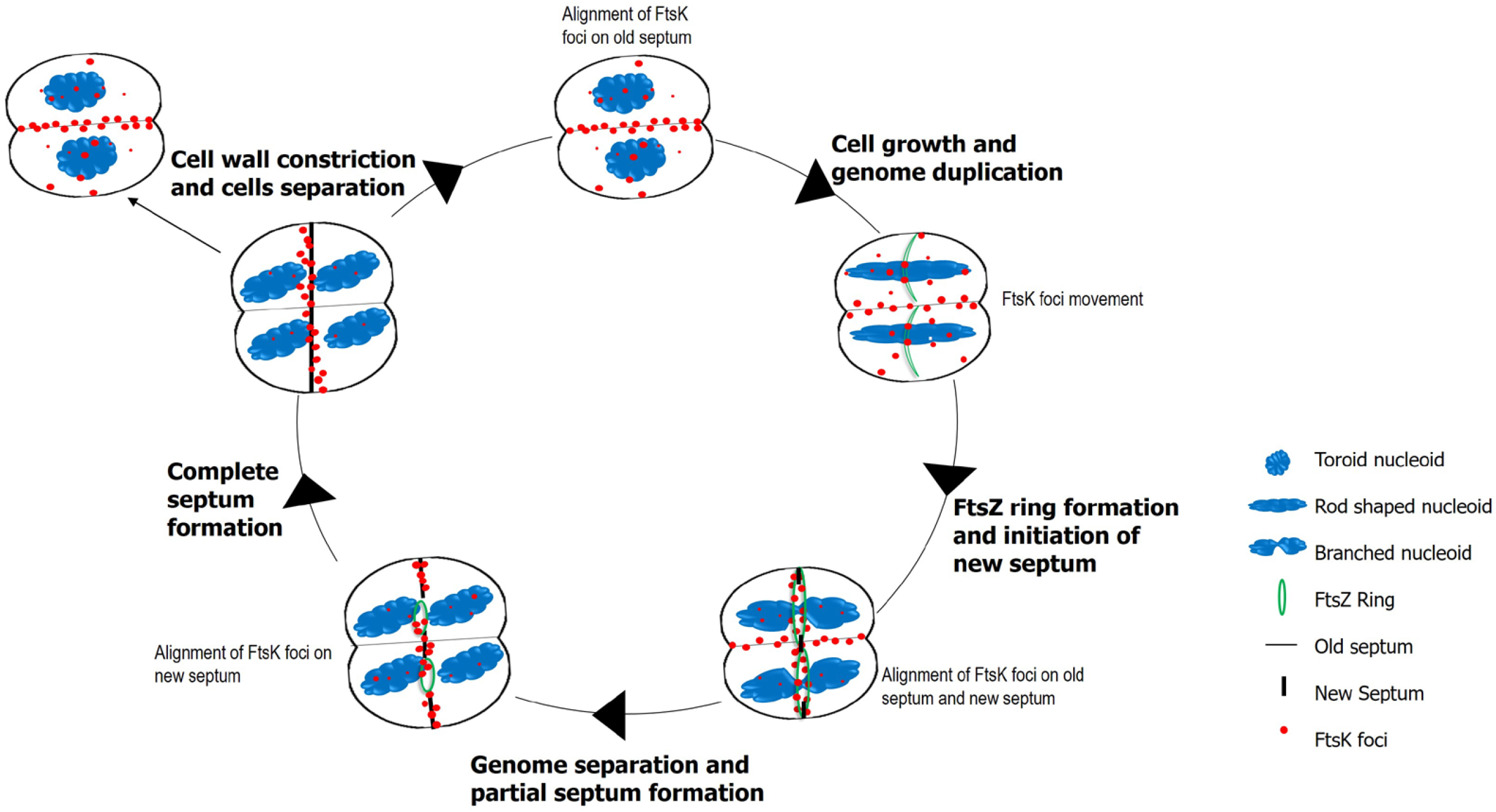
Schematic model of the dynamics of FtsK in *D. radiodurans* during cell division. Based on the biochemical assays and cellular localization studies, the probable role and dynamics of FtsK protein in *D. radiodurans* were proposed. FtsK forms foci on nucleoid as well as membrane. Initially, a maximum number of FtsK foci are aligned at the septum. As the cell grows and the genome duplicates, FtsK foci move. With the formation of the FtsZ ring and subsequent initiation of the new septa, the alignment of FtsK shifts perpendicularly from the old one to the new one. When new septum formation completes, FtsK moves entirely to the new septa before any cell wall constriction starts and the whole process repeats in the next cell division. The ATPase activity, sequence-specific DNA binding activity, and cellular interaction of FtsK with other segrosome and divisome proteins may aid in the smooth and coordinated progression of different cellular processes to maintain the genome stability in *D. radiodurans*.

FtsK/SpoIIIE DNA translocases are multidomain and multifunctional proteins that are part of bacterial cell divisome machinery (Mohone and Goley, 2020). In *D. radiodurans,* our earlier work showed the involvement of ParBs, TopoIB in genome segregation, and FtsZ, FtsA, and DivIVA in cell division (Maurya et al. 2019, Modi et al. 2014, Chaudhary et al. 2019, Kota et al. 2021). The co-immunoprecipitation results showed that drFtsK interacts with these proteins indicating its role in both these processes. When different domain-specific *drftsK* deletion mutants were generated, the growth rate of mutants was significantly reduced during the early and exponential phases. Remarkably, mutants showed a high growth rate during the stationary phase. Microscopy data showed improper membrane staining and membrane bulges in *ftsK* mutants. Membrane bulges were also observed earlier in *S. aureus ΔftsK/ΔspoIIIE* mutants (Veiha et al., 2017). This indicates the role of drFtsK in envelope reorganization during cell division and in mutants, the lack of this function might have caused the delay in the growth rate. (Geissler and Margolin, 2005). Previously, the movement of DNA through the tetrad compartments during PIR was shown to be necessary for the radiation resistance phenotype (Levin-Zaidman et al. 2003). The reduced growth rate in mutants exposed to gamma radiation could also be attributed to the delayed DNA translocation through the septum. Further, the fast recovery of the mutants at the stationary phase can be speculated to be due to the bypass of FtsK, as seen earlier in *E.coli* by FtsA and FtsN interaction (Pichoff et al., 2018).

The genome segregation process in *E. coli* progresses from the *ori*-to-*ter* loci and FtsK protein was actively involved in the “ter” locus positioning at mid cell region (Stouf et. al., 2013). In *Vibrio cholerae,* two divergent *difs*, chromosome resolution sites were identified for two chromosomes, and both are activated by FtsK (Val et al. 2008). In *D. radiodurans* also, genome sequence analysis has revealed putative “*dif”* sites and six putative tyrosine recombinases on both primary and secondary chromosomes (Figure S2). An independent study will be conducted to check the functional interacting partners of drFtsK among these recombinases and their role in chromosome dimer resolution at “*dif”* sites.

Recent electron cryo-microscopy (cryo-EM) structure studies of dsDNA bound FtsK_αβ_ from *P. aeruginosa* revealed changes in different conformational states of ATPase domains in homo-hexameric rings generate the translocation movement on DNA (Jean et al., 2020). drFtsK also forms homo-hexamers and the time-lapse microscopy results showed that FtsK-RFP is dynamic, orientating its translocating activity along the new septum from the old septum in dividing cells. Based on the observed results, FtsK movement can be linked to the cellular growth stage where the active role of protein in the smooth coordination of genome segregation and cell division processes can be thought of (Figure 10). As shown in the pictorial form, when the cell divides into two daughter cells, most of the FtsK aligns on the old septum. During the subsequent cell growth, at the time of genome duplication and elongation, FtsK disperses on the genome. At the early cell division stage of FtsZ ring formation and initial constriction appearance, FtsK alignment shifts from the old to the new septum. Foci alignment on the new septum is more prominent at the later stages of genome segregation and cell septum formation. Thus, FtsK marks the new septum and the cell divides into two after complete septum formation and cell wall constriction. All the results suggest that the dynamic multiprotein interactions coordinate accurate segregation of duplicated intertwined circular genome elements as well as the next plane of cell division in this coccus. In conclusion, the drFtsK characterized in this report is a multifunctional protein having ATPase, and DNA binding activities and has a role in the determination of cell architecture, in general, and a faithful inheritance of its multipartite genome system.

## Acknowledgments

We acknowledge Dr. **Ganesh K. Maurya** and Dr. **Reema Chaudhary** for their help in *in-silico* studies and co-immunoprecipitation experiments. We also thank Prof. **Ian Grainge** for providing materials for conducting the site-specific recombination assay.

## Authors’ contributions

**Shruti Mishra** proposed the functions of the protein, conducted the experiments, analyzed the data, and wrote the manuscript. **Hari S Misra** conceived the idea, analyzed the data, wrote the manuscript, and is the principal investigator. **Swathi Kota** conceived the idea, designed the experiments, analyzed the data, wrote the manuscript, and communicated for publication.

---

**Figure S1:** Multiple sequence alignment of the full-length sequences of FtsK/SpoIIIE protein family members across different species-*Deinococcus* r*adiodurans* (FtsK_Dr), SpoIIIE from *B. subtilis* (SpoIIIE_Bs), FtsK of *Escherichia coli* (FtsK_Ec), *Pseudomonas aeruginosa* (FtsK_Pa)*, Lactococcus lactis* (FtsK_Ll)*, Vibrio cholerae* (FtsK_Vc), and *Staphylococcus aureus* (FtsK_Sa) using PROMALS3D webserver. Predicted secondary structures are displayed below the sequences Boundaries of the conserved motifs in the C-terminal are marked as a black box for walker A domain (ATP binding and ATPase activity), a purple box for walker B motif, and a red box for the DNA binding motif (A). The neighbor-joining phylogenetic tree (without distance corrections) of FtsK/SpoIIIE family proteins across different species (mentioned above) was constructed using the PHYLIP program (B).

**Figure S2:** Putative tyrosine recombinases annotated in the genome of *D. radiodurans*. (A) Bioinformatics analysis was done using *E. coli* XerC and XerD sequences as templates to identify putative tyrosine recombinases in the *D. radiodurans.* A total of six putative ORFs were identified with XerCD-specific domains which could function as tyrosine recombinases (B).

**Figure S3:** The oligomeric status and ATPase activity of drFtsK (FtsKΔN). The oligomeric status of the protein was checked by dynamic light scattering in the presence/absence of *E. coli* KOPS containing DNA and ATP. FtsKΔN exists as pre-formed hexamers with a peak at ∼21 d.nm (A). ATPase assay of the protein was performed in the presence of *E. coli* KOPS containing dsDNA. Different concentrations of protein (0.1, 0.2, 0.3, and 0.5 µM) were incubated with assay buffer with and without DNA as described in the methodology and ATPase activity was calculated (B).

**Figure S4:** Distribution of KOPS-GGGCAGGG (A) and GGGNAGGG (B) in chromosome I and chromosome II of *D. radiodurans* and *Escherichia coli*. The distribution of 8 mer KOPS on the lagging strand and leading strand is analyzed by the DistAMo tool and shown separately. A significant over-or under-representation is colour-coded by a red or blue colour, respectively. Rings from outside to the inside differ in the size of the sliding window from 50 to 500kb in the 50kb step.

**Figure S5:** PAGE gels depicting the purified recombinant deinococcal FtsK gamma domain (drFtsKγ), *E. coli* XerC (ecXerC), *E. coli* XerD (ecXerD), and *E. coli* FtsK gamma domain (ecFtsKγ) proteins from transgenic *E. coli* BL 21 strains.

**Figure S6:** Confirmation of the insertional replacement of different regions of the coding sequence of *ftsK*. Schematic representation of the domain organization in drFtsK showing the regions deleted to generate deletion mutants as described in methods; Δ*ftsK*-full-length FtsK deletion mutant*, ΔftsKMC-* FtsK middle and C-terminal deletion mutant, and *ΔftsKN-* FtsK N-terminal deletion mutant (A). Diagnostic PCR was done to check the successful replacement of different regions’ coding sequence of *ftsK* with antibiotic cassette (*npt*II) by using sequence-specific primers for *nptII*-NF and NR, *ftsKN*-AF and AR, *ftsKC* – CF and CR, *nptII –FtsKM*-NF and MR (B) and flanking primers-UF and DR (C). Primer details are mentioned in Table S2.

**Figure S7:** Effect of different domain deletion of *drftsK* on the phenotype of *D. radiodurans* cells during exponential phase at 6 hrs (A) and 9 hrs (B). FtsK mutants formed membrane bulges and the percent of cells showing this phenotype were counted and plotted at both 6 hrs and 9 hrs (C).

**Table S1:** List of all the strains and plasmids used in this study.

**Table S2:** List of primers used in this study.

## References

1. Aarsman, M. E., Piette, A., Fraipont, C., Vinkenvleugel, T. M., Nguyen-Distèche, M., & den Blaauwen, T. (2005). Maturation of the Escherichia coli divisome occurs in two steps. Molecular microbiology, 55(6), 1631–1645.

2. Aussel, L., Barre, F. X., Aroyo, M., Stasiak, A., Stasiak, A. Z., & Sherratt, D. (2002). FtsK is a DNA motor protein that activates chromosome dimer resolution by switching the catalytic state of the XerC and XerD recombinases. Cell, 108(2), 195–205.

3. Becker, E. C., & Pogliano, K. (2007). Cell-specific SpoIIIE assembly and DNA translocation polarity are dictated by chromosome orientation. Molecular microbiology, 66(5), 1066–1079.

4. Begg, K. J., Dewar, S. J., & Donachie, W. D. (1995). A new Escherichia coli cell division gene, ftsK. Journal of bacteriology, 177(21), 6211–6222.

5. Berezuk, A. M., Glavota, S., Roach, E. J., Goodyear, M. C., Krieger, J. R., & Khursigara, C. M. (2018). Outer membrane lipoprotein RlpA is a novel periplasmic interaction partner of the cell division protein FtsK in Escherichia coli. Scientific reports, 8(1), 1–14.

6. Berezuk, A. M., Goodyear, M., & Khursigara, C. M. (2014). Site-directed fluorescence labeling reveals a revised N-terminal membrane topology and functional periplasmic residues in the *Escherichia coli* cell division protein FtsK. Journal of Biological Chemistry, 289(34), 23287–23301.

7. Bigot, S., Sivanathan, V., Possoz, C., Barre, F. X., & Cornet, F. (2007). FtsK, a literate chromosome segregation machine. Molecular microbiology, 64(6), 1434–1441.

8. Bisicchia, P., Steel, B., Mariam Debela, M. H., Löwe, J., & Sherratt, D. (2013). The N-terminal membrane-spanning domain of the Escherichia coli DNA translocase FtsK hexamerizes at midcell. MBio, 4(6), e00800–13.

9. Cattoni, D. I., Chara, O., Godefroy, C., Margeat, E., Trigueros, S., Milhiet, P. E., & Nöllmann, M. (2013). SpoIIIE mechanism of directional translocation involves target search coupled to sequence-dependent motor stimulation. EMBO reports, 14(5), 473–479.

10. Charaka, V. K., & Misra, H. S. (2012). Functional characterization of the role of the chromosome I partitioning system in genome segregation in Deinococcus radiodurans. Journal of bacteriology, 194(21), 5739–5748.

11. Chaudhary, R., Gupta, A., Kota, S., & Misra, H. S. (2019). N-terminal domain of DivIVA contributes to its dimerization and interaction with genome segregation proteins in a radioresistant bacterium *Deinococcus radiodurans*. International journal of biological macromolecules, 128, 12–21

12. Chaudhary, R., Kota, S., & Misra, H. S. (2021). DivIVA Regulates Its Expression and the Orientation of New Septum Growth in *Deinococcus radiodurans*. Journal of bacteriology, 203(15), e00163–21.

13. Cox M.M, Battista J.R. (2005). *Deinococcus radiodurans* - the consummate survivor. Nat Rev Microbiol. Nov;3(11):882–92. doi: 10.1038/nrmicro1264. PMID: 16261171.

14. den Blaauwen, T., Hamoen, L. W., & Levin, P. A. (2017). The divisome at 25: the road ahead. Current opinion in microbiology, 36, 85–94.

15. Di Lallo, G., Fagioli, M., Barionovi, D., Ghelardini, P., & Paolozzi, L. (2003). Use of a two-hybrid assay to study the assembly of a complex multicomponent protein machinery: bacterial septosome differentiation. Microbiology, 149(12), 3353–3359.

16. Draper, G. C., McLennan, N., Begg, K., Masters, M., & Donachie, W. D. (1998). Only the N-terminal domain of FtsK functions in cell division. Journal of bacteriology, 180(17), 4621–4627.

17. Dubarry, N., Possoz, C., & Barre, F. X. (2010). Multiple regions along the Escherichia coli FtsK protein are implicated in cell division. Molecular microbiology, 78(5), 1088–1100.

18. Errington, J., Bath, J., & Wu, L. J. (2001). DNA transport in bacteria. Nature Reviews Molecular Cell Biology, 2(7), 538–545.

19. Floc’h, K., Lacroix, F., Servant, P., Wong, Y. S., Kleman, J. P., Bourgeois, D., & Timmins, J. (2019). Cell morphology and nucleoid dynamics in dividing Deinococcus radiodurans. Nature communications, 10(1), 1–13.

20. Geissler, B., & Margolin, W. (2005). Evidence for functional overlap among multiple bacterial cell division proteins: compensating for the loss of FtsK. Molecular microbiology, 58(2), 596–612.

21. Goehring, N. W., & Beckwith, J. (2005). Diverse paths to midcell: assembly of the bacterial cell division machinery. Current biology, 15(13), R514–R526.

22. Goehring, N. W., Gueiros-Filho, F., & Beckwith, J. (2005). Premature targeting of a cell division protein to midcell allows dissection of divisome assembly in Escherichia coli. Genes & development, 19(1), 127–137.

23. Goehring, N. W., Petrovska, I., Boyd, D., & Beckwith, J. (2007). Mutants, suppressors, and wrinkled colonies: mutant alleles of the cell division gene ftsQ point to functional domains in FtsQ and a role for domain 1C of FtsA in divisome assembly. Journal of bacteriology, 189(2), 633–645.

24. Grainge, I. (2010). FtsK–a bacterial cell division checkpoint?. Molecular microbiology, 78(5), 1055–1057.

25. Grainge, I., Lesterlin, C., & Sherratt, D. J. (2011). Activation of XerCD-dif recombination by the FtsK DNA translocase. Nucleic acids research, 39(12), 5140–5148.

26. Green, M. R., & Sambrook, J. (2012). A laboratory manual. John Inglis, (2012).

27. Ip, S. C., Bregu, M., Barre, F. X., & Sherratt, D. J. (2003). Decatenation of DNA circles by FtsK-dependent Xer site-specific recombination. The EMBO journal, 22(23), 6399–6407.

28. Jean, N. L., Rutherford, T. J., & Löwe, J. (2020). FtsK in motion reveals its mechanism for double-stranded DNA translocation. Proceedings of the National Academy of Sciences, 117(25), 14202–14208.

29. Karimova, G., Pidoux, J., Ullmann, A., & Ladant, D. (1998). A bacterial two-hybrid system based on a reconstituted signal transduction pathway. Proceedings of the National Academy of Sciences USA, 95(10), 5752–5756.

30. Keller, A. N., Xin, Y., Boer, S., Reinhardt, J., Baker, R., Arciszewska, L. K., … & Grainge, I. (2016). Activation of Xer-recombination at dif: structural basis of the FtsKγ–XerD interaction. Scientific reports, 6(1), 1–12.

31. Khairnar, N. P., Kamble, V. A., & Misra, H. S. (2008). RecBC enzyme overproduction affects UV and gamma radiation survival of *Deinococcus radiodurans*. DNA repair, 7(1), 40–47.

32. Kota, S., & Misra, H. S. (2006). PprA: a protein implicated in radioresistance of Deinococcus radiodurans stimulates catalase activity in Escherichia coli. Applied microbiology and biotechnology, 72(4), 790–796.

33. Kota, S., Chaudhary, R., Mishra, S., & Misra, H. S. (2021). Topoisomerase IB interacts with genome segregation proteins and is involved in multipartite genome maintenance in *Deinococcus radiodurans*. Microbiological Research, 242, 126609

34. Leonard, T. A., Butler, P. J., & Löwe, J. (2005). Bacterial chromosome segregation: structure and DNA binding of the Soj dimer—a conserved biological switch. The EMBO journal, 24(2), 270–282.

35. Lesterlin, C., Mercier, R., Boccard, F., Barre, F. X., & Cornet, F. (2005). Roles for replichores and macrodomains in segregation of the *Escherichia coli* chromosome. EMBO reports, 6(6), 557–562.

36. Levin-Zaidman, S., Englander, J., Shimoni, E., Sharma, A. K., Minton, K. W., & Minsky, A. (2003). Ringlike structure of the *Deinococcus radiodurans* genome: a key to radioresistance? Science, 299(5604), 254–256.

37. Levy, O., Ptacin, J. L., Pease, P. J., Gore, J., Eisen, M. B., Bustamante, C., & Cozzarelli, N. R. (2005). Identification of oligonucleotide sequences that direct the movement of the Escherichia coli FtsK translocase. Proceedings of the National Academy of Sciences, 102(49), 17618–17623.

38. Löwe, J., Ellonen, A., Allen, M. D., Atkinson, C., Sherratt, D. J., & Grainge, I. (2008). Molecular mechanism of sequence-directed DNA loading and translocation by FtsK. Molecular cell, 31(4), 498–509.

39. Mahone, C. R., & Goley, E. D. (2020). Bacterial cell division at a glance. Journal of Cell Science, 133(7), jcs237057.

40. Maier, R.M. 2009. Bacterial growth, p 37–54. In Maier RM, Pepper IL, Gerb CP (ed), Environmental toxicology, part I: review of basic microbiological concepts, 2nd ed. Elsevier Academic Press, San Diego, CA.

41. Makarova K.S., Aravind L., Wolf Y.I., Tatusov R.L., Minton K.W., Koonin E.V., Daly M.J. (2001). Genome of the extremely radiation-resistant bacterium Deinococcus radiodurans viewed from the perspective of comparative genomics. Microbiol Mol Biol Rev. Mar;65(1):44–79. doi: 10.1128/MMBR.65.1.44-79.2001. PMID: 11238985; PMCID: PMC99018.

42. Maurya, G. K., Kota, S., & Misra, H. S. (2019). Characterisation of ParB encoded on multipartite genome in Deinococcus radiodurans and their roles in radioresistance. Microbiological research, 223, 22–32.

43. Maurya, G. K., Modi, K., & Misra, H. S. (2016). Divisome and segrosome components of Deinococcus radiodurans interact through cell division regulatory proteins. Microbiology, 162(8), 1321–1334.

44. Maurya, G. K., Modi, K., Banerjee, M., Chaudhary, R., Rajpurohit, Y. S., & Misra, H. S. (2018). Phosphorylation of FtsZ and FtsA by a DNA damage-responsive Ser/Thr protein kinase affects their functional interactions in Deinococcus radiodurans. Msphere, 3(4), e00325–18.

45. Midonet, C., Das, B., Paly, E., & Barre, F. X. (2014). XerD-mediated FtsK-independent integration of TLCϕ into the *Vibrio cholerae* genome. Proceedings of the National Academy of Sciences, 111(47), 16848–16853.

46. Mishra, S., Chaudhary, R., Singh, S., Kota, S., & Misra, H. S. (2019). Guanine quadruplex DNA regulates gamma radiation response of genome functions in the radioresistant bacterium *Deinococcus radiodurans*. Journal of Bacteriology, 201(17), e00154–19.

47. Misra, H. S., Khairnar, N. P., Kota, S., Shrivastava, S., Joshi, V. P., & Apte, S. K. (2006). An exonuclease I-sensitive DNA repair pathway in *Deinococcus radiodurans*: a major determinant of radiation resistance. Molecular microbiology, 59(4), 1308–1316.

48. Misra, H. S., Rajpurohit, Y. S., & Kota, S. (2013). Physiological and molecular basis of extreme radioresistance in *Deinococcus radiodurans*. Current Science(Bangalore), 104(2), 194–205.

49. Modi, K., & Misra, H. S. (2014). Dr-FtsA, an actin homologue in *Deinococcus radiodurans* differentially affects Dr-FtsZ and Ec-FtsZ functions in vitro. PloS one, 9(12), e115918.

50. Pazos, M., Natale, P., & Vicente, M. (2013). A specific role for the ZipA protein in cell division: stabilization of the FtsZ protein. Journal of Biological Chemistry, 288(5), 3219–3226.

51. Pei, J., Kim, B. H., & Grishin, N. V. (2008). PROMALS3D: a tool for multiple protein sequence and structure alignments. Nucleic acids research, 36(7), 2295–2300.

52. Pichoff, S., Du, S., & Lutkenhaus, J. (2018). Disruption of divisome assembly rescued by FtsN–FtsA interaction in Escherichia coli. Proceedings of the National Academy of Sciences, 115(29), E6855–E6862.

53. Repar, J., Zahradka, D., Sović, I., & Zahradka, K. (2021). Characterization of gross genome rearrangements in Deinococcus radiodurans recA mutants. Scientific reports, 11(1), 1–11.

54. Schäfer, M., Schmitz, C., Facius, R., Horneck, G., Milow, B., Funken, K. H., & Ortner, J. (2000). Systematic study of parameters influencing the action of rose bengal with visible light on bacterial cells: comparison between the biological effect and singlet-oxygen production. Photochemistry and photobiology, 71(5), 514–523.

55. Slade, D., & Radman, M. (2011). Oxidative stress resistance in *Deinococcus radiodurans*. Microbiology and molecular biology reviews, 75(1), 133–191.

56. Sobetzko, P., Jelonek, L., Strickert, M., Han, W., Goesmann, A., & Waldminghaus, T. (2016). DistAMo: a web-based tool to characterize DNA-motif distribution on bacterial chromosomes. Frontiers in microbiology, 7, 283.

57. Steiner, W. W., & Kuempel, P. L. (1998). Sister chromatid exchange frequencies in Escherichia coli analyzed by recombination at the dif resolvase site. Journal of Bacteriology, 180(23), 6269–6275.

58. Stouf, M., Meile, J. C., & Cornet, F. (2013). FtsK actively segregates sister chromosomes in *Escherichia coli*. Proceedings of the National Academy of Sciences, 110(27), 11157–11162.

59. Tiyanont, K., Doan, T., Lazarus, M. B., Fang, X., Rudner, D. Z., & Walker, S. (2006). Imaging peptidoglycan biosynthesis in Bacillus subtilis with fluorescent antibiotics. Proceedings of the National Academy of Sciences, 103(29), 11033–11038.

60. Val, M. E., Kennedy, S. P., El Karoui, M., Bonne, L., Chevalier, F., & Barre, F. X. (2008). FtsK-dependent dimer resolution on multiple chromosomes in the pathogen Vibrio cholerae. PLoS genetics, 4(9), e1000201.

61. Veiga, H., & G. Pinho, M. (2017). S taphylococcus aureus requires at least one F ts K/S po IIIE protein for correct chromosome segregation. Molecular Microbiology, 103(3), 504–517.

62. White, O., Eisen, J. A., Heidelberg, J. F., Hickey, E. K., Peterson, J. D., Dodson, R. J.,… & Fraser, C. M. (1999). Genome sequence of the radioresistant bacterium *Deinococcus radiodurans* R1. Science, 286(5444), 1571–1577.

63. Yates, J., Aroyo, M., Sherratt, D. J., & Barre, F. X. (2003). Species specificity in the activation of Xer recombination at dif by FtsK. Molecular microbiology, 49(1), 241–249.

64. Yu, X. C., Tran, A. H., Sun, Q., & Margolin, W. (1998). Localization of cell division protein FtsK to the *Escherichia coli* septum and identification of a potential N-terminal targeting domain. Journal of Bacteriology, 180(5), 1296–1304.

