## Supplementary material for "FtsK, a DNA motor protein, coordinates the genome segregation and early cell division processes in *Deinococcus radiodurans*"

(A)

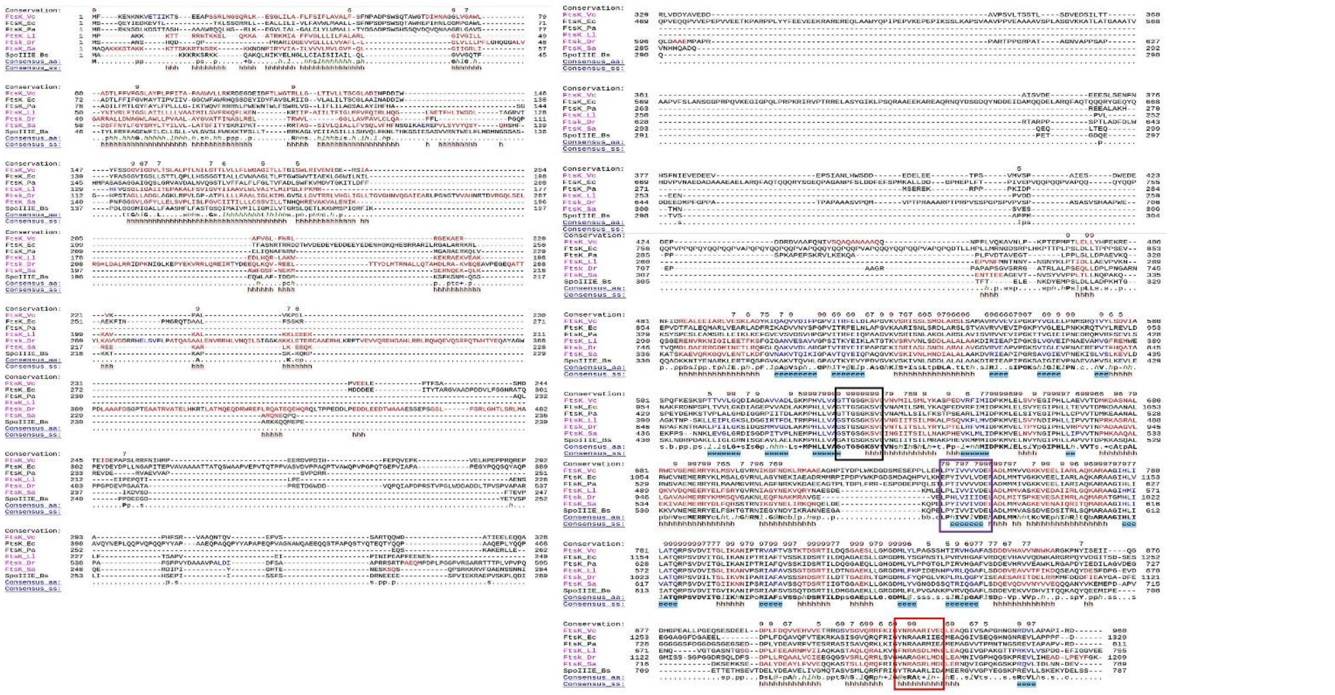

(B)

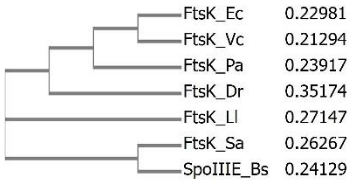


**Figure S1:** Multiple sequence alignment of the full-length sequences of FtsK/SpoIIIE protein family members across different species- *Deinococcus radiodurans* (FtsK_Dr), SpoIIIE from *B. subtilis* (SpoIIIE_Bs), FtsK of *Escherichia coli* (FtsK_Ec), *Pseudomonas aeruginosa* (FtsK_Pa), *Lactococcus lactis* (FtsK_Ll), *Vibrio cholerae* (FtsK_Vc), and *Staphylococcus aureus* (FtsK_Sa) using PROMALS3D webserver. Predicted secondary structures are displayed below the sequences Boundaries of the conserved motifs in the C-terminal are marked as a black box for walker A domain (ATP binding and ATPase activity), a purple box for walker B motif, and a red box for the DNA binding motif (A). The neighbour-joining phylogenetic tree (without distance corrections) of FtsK/SpoIIIE family proteins across different species (mentioned above) was constructed using the PHYLIP program (B).

(A)

| Gene locus tag | Location | Annotation |
|---|---|---|
| E5E91_RS02620 | Chr I | Site specific integrase |
| E5E91_RS13705 | Chr II | DDE-type integrase/transposase/recombinase |
| E5E91_RS14115 | Chr II | Site specific integrase |
| E5E91_RS14250 | Chr II | Site specific integrase |
| E5E91_RS15640 | Megaplasmid | Tyrosine-type recombinase/integrase |
| E5E91_RS15930 | Small plasmid | Site specific integrase |

(B)

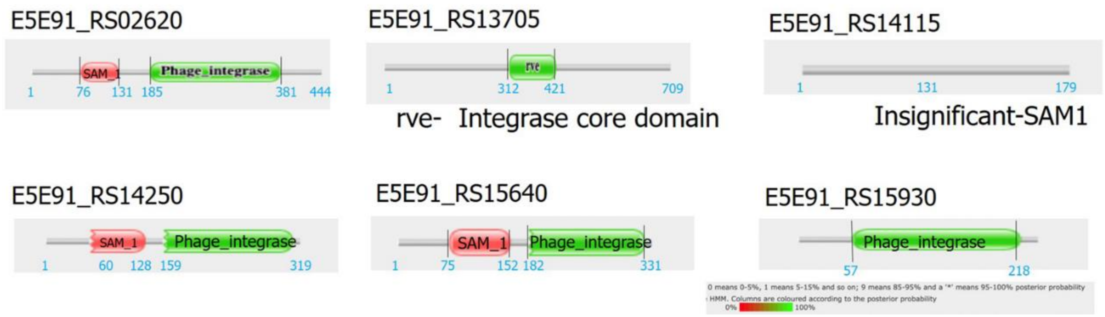


**Figure S2**: Putative tyrosine recombinases annotated in the genome of *D. radiodurans*. (A) Bioinformatics analysis was done using *E. coli* XerC and XerD sequences as templates to identify putative tyrosine recombinases in the *D. radiodurans*. A total of six putative ORFs were identified with XerCD-specific domains which could function as tyrosine recombinases (B).

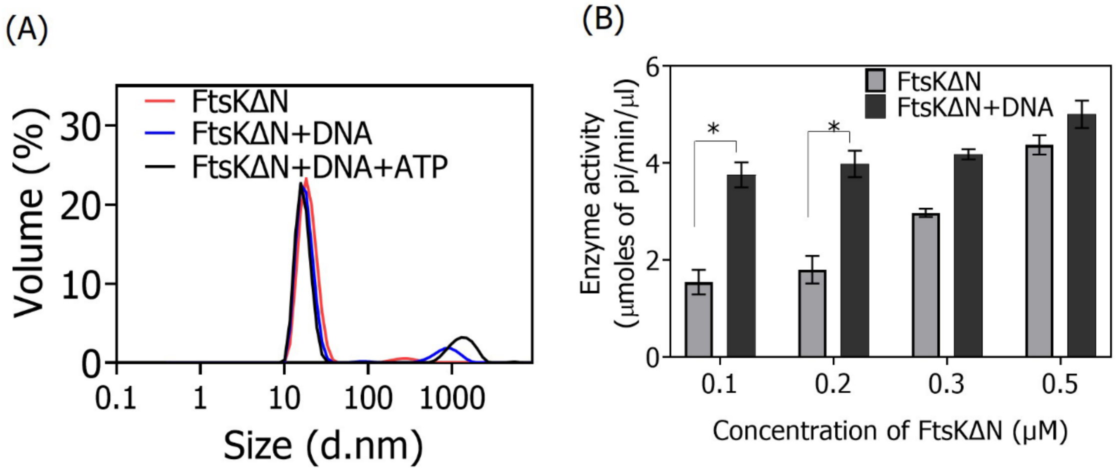


**Figure S3**: The oligomeric status and ATPase activity of drFtsK (FtsKΔN). The oligomeric status of the protein was checked by dynamic light scattering in the presence/absence of *E. coli* KOPS containing DNA and ATP. FtsKΔN exists as pre-formed hexamers with peak at ~21 d.nm (A). ATPase assay of the protein was performed in the presence of *E. coli* KOPS containing dsDNA. Different concentrations of protein (0.1, 0.2, 0.3 and 0.5 μM) were incubated with assay buffer with and without DNA as described in the methodology and ATPase activity was calculated (B).

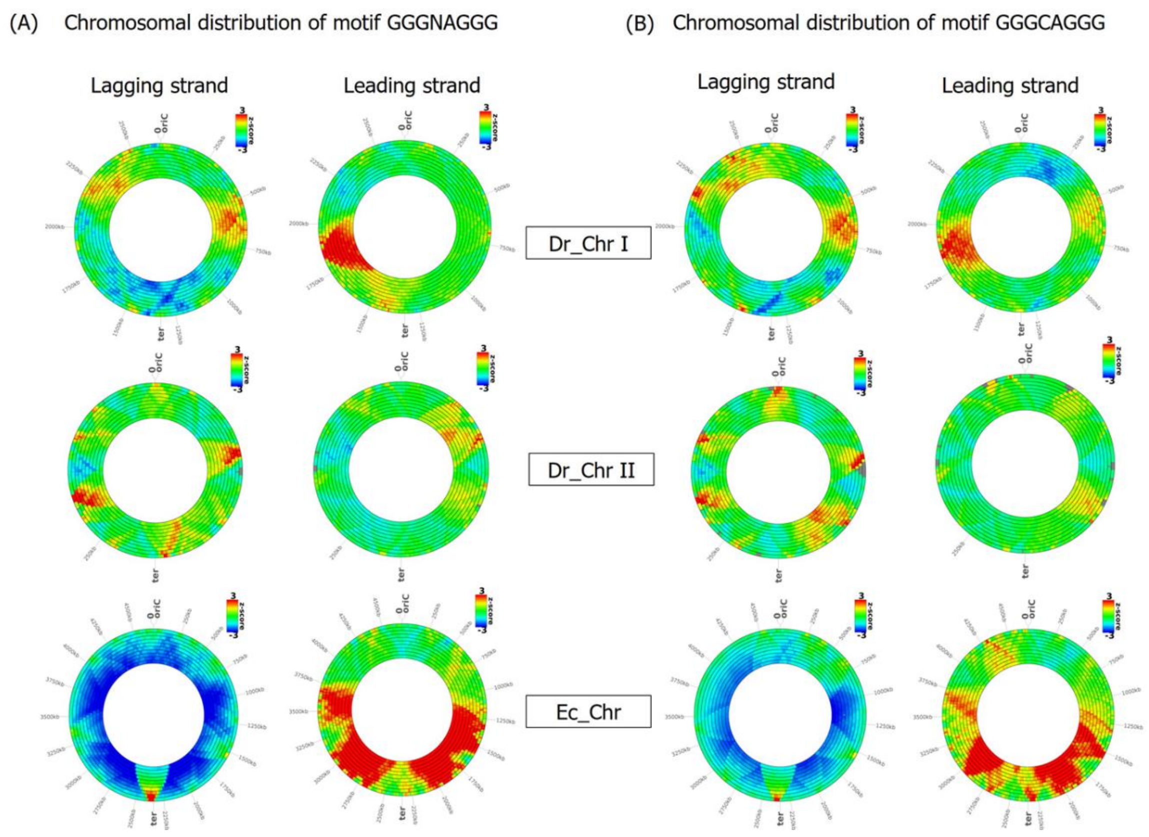


**Figure S4:** Distribution of KOPS- GGGCAGGG (A) and GGGNAGGG (B) in chromosome I and chromosome II of *D. radiodurans* and *Escherichia coli*. The distribution of 8 mer KOPS on the lagging strand and leading strand is analyzed by the DistAMo tool and shown separately. A significant over-or under-representation is colour-coded by a red or blue colour, respectively. Rings from outside to the inside differ in the size of the sliding window from 50 to 500kb in the 50kb step.

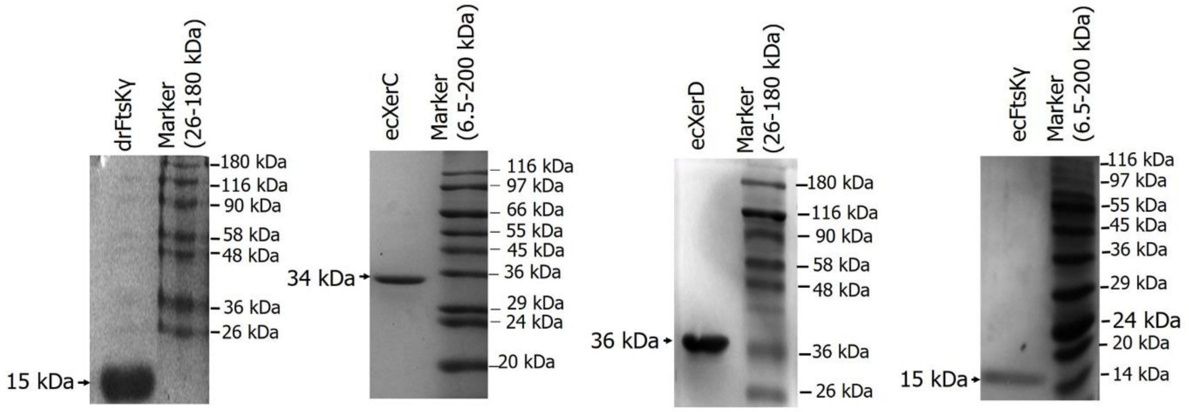



**Figure S5:** PAGE gels depicting the purified recombinant deinococcal FtsK gamma domain (drFtsKγ), *E. coli* XerC (ecXerC), *E. coli* XerD (ecXerD) and *E. coli* FtsK gamma domain (ecFtsKγ) proteins from transgenic *E. coli* BL 21 strains.

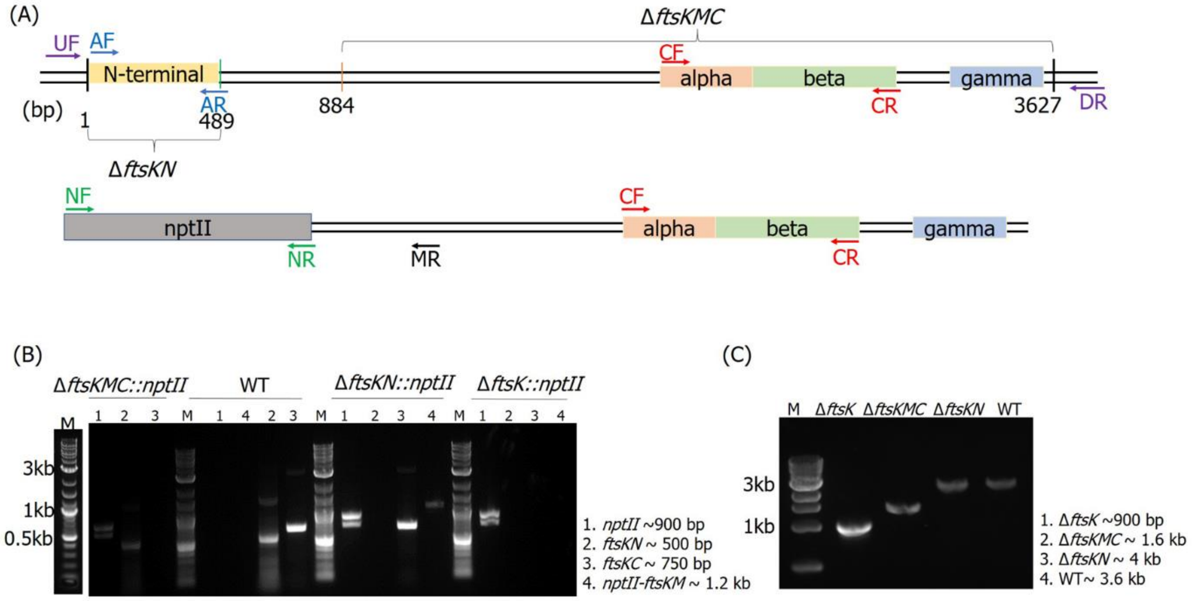



**Figure S6:** Confirmation of the insertional replacement of different regions of the coding sequence of *ftsK*. Schematic representation of the domain organization in drFtsK showing the regions deleted to generate deletion mutants as described in methods; Δ*ftsK*- full-length FtsK deletion mutant, *ΔftsKMC*- FtsK middle and C-terminal deletion mutant and *ΔftsKN*- FtsK N-terminal deletion mutant (A). Diagnostic PCR was done to check the successful replacement of different regions' coding sequence of *ftsK* with antibiotic cassette (*npt*II) by using sequence-specific primers for *nptII*- NF and NR, *ftsKN*- AF and AR, *ftsKC* – CF and CR, *nptII –FtsKM*- NF and MR (B) and flanking primers-UF and DR (C). Primer details are mentioned in the Table S2.

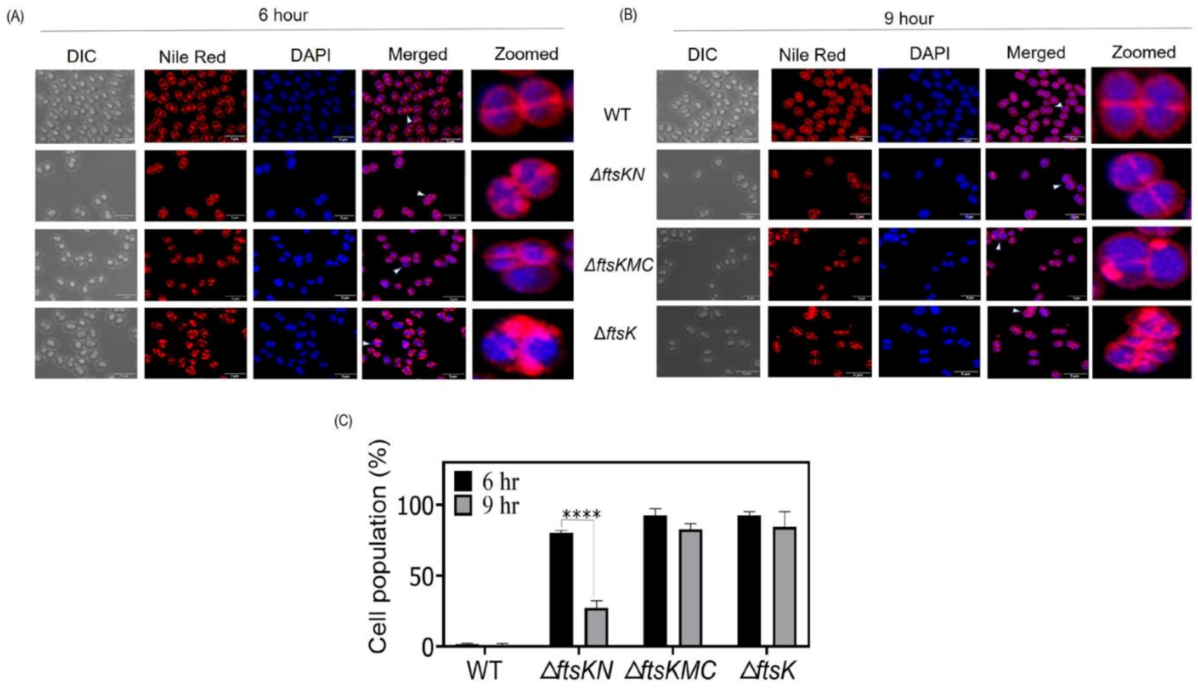


**Figure S7:** Effect of different domain deletion of *drftsK* on the phenotype of *D. radiodurans* cells during exponential phase at 6 hrs (A) and 9 hrs (B). FtsK mutants formed membrane bulges and the percent of cells showing this phenotype were counted and plotted at both 6 hrs and 9 hrs (C).

Table S1: List of all the strains and plasmids used in this study.

| S.No. | Strain | Genotype | Source |
|---|---|---|---|
| 1 | *D. radiodurans* R1 | wild type | lab stock |
| 2 | *E. coli* MG1655 | Wild type | lab stock |
| 3 | *E. coli* DH5α | *F- / endA1 hsdR17 glnV44 thi-1 recA1 gyrArelA* Δ *(lacIZYA-argF) U169 deoR (Φ80dlac*Δ *(lacZ)M15)* | lab stock |
| 4 | *E.coli* BL21 (DE3) *pLys*S | *F-ompT gal [dcm] [lon] hsdSB DE3::T7RNA* | lab stock |
| 5 | Δ*ftsK* | FtsK knock-out mutant ($Kan^R$, 8µg/ml) | This study |
| 6 | *ΔftsKMC* | Middle and C-terminal of FtsK deletion mutant ($Kan^R$, 8µg/ml) | This study |
| 7 | *ΔftsKN* | N-terminal of FtsK deletion mutant ($Kan^R$, 8µg/ml) | This study |
| 8 | *ftsK::ftsK-rfp* | *ftsK* replaced with *ftsK-rfp* under its native promoter ($Kan^R$, 8µg/ml) | This study |
| 9 | Δ*ftsK*+FtsZ-GFP | Knock-out mutant of *ftsK* harbouring pVHZGFP (KanR, 8µg/ml + $Spec^R$, 70µg/ml ) | This study |
| 10 | Dr+FtsZ-GFP | D. radiodurans R1 harbouring pVHZGFP ($Kan^R$, 8µg/ml + $Spec^R$, 70µg/ml ) | Modi et al., 2014 |
| 11 | *ftsK::ftsK-rfp*+FtsZ-GFP | Knock-in mutant of *ftsK-rfp* harbouring pVHZGFP ($Kan^R$,8µg/ml + $Spec^R$,70µg/ml) | This study |

| S.No. | Plasmids | Description | Source |
|---|---|---|---|
| 1 | pET28a(+) | ~ 5.3 kb plasmid; N-terminal 6XHis tag , $Kan^R$ | Novagen |
| 2 | pDsRed | ~ 3.3 kb plasmid; N-terminal red fluorescent tag, $Amp^R$ | Clontech Inc. |
| 3 | pNOKOUT | $pBSK^+$ ($Amp^R$) containing *nptII*cassette (937bp), ~4.0 kb, $Amp^R$/ $Kan^R$ | Khairnar et al., 2008 |

| | | | |
|---|---|---|---|
| 4 | pUT18 | pUC19 + T18 fragment of CyaA under *lac* promoter. ~3.0 kb, $Amp^R$ | Karimova et al., 2005 |
| 5 | pET0400 | pET-28a + Dr_0400 (ORF), ~8.2kb, $Kan^R$ | This study |
| 6 | pETFtsK | pET-28a + *ftsK* (Dr_0400 + Dr_0401), ~8.9kb, $Kan^R$ | This study |
| 7 | pETDFKγ | pET-28a + *ecftsk-γ*, ~5.7kb, $Kan^R$ | This study |
| 8 | pETEFKγ | pET-28a + *drftsk-γ,* ~5.7kb, $Kan^R$ | This study |
| 9 | pETEXC | pET-28a + *ecxerc*, ~6.2 kb, $Kan^R$ | This study |
| 10 | pETEXD | pET-28a + *ecxerd*, ~6.3 kb, $Kan^R$ | This study |
| 11 | pUTDFZ | pUT18 carrying *drftsZ*, ~4 kb, $Amp^R$ | Modi & Misra, 2014 |
| 12 | pUTDFA | pUT18 carrying *drftsA,*~4.4 kb, $Amp^R$ | Modi & Misra, 2014 |
| 13 | pUTB2 | pUT18 carrying *drparB2*, ~3.9kb, $Amp^R$ | Maurya et al., 2016 |
| 14 | pUTB3 | pUT18 carrying *drparB3*, ~3.9kb, $Amp^R$ | Maurya et al., 2016 |
| 15 | pUTB4 | pUT18 carrying *drparB4*, ~3.9kb, $Amp^R$ | Maurya et al., 2016 |
| 16 | pUTTopoIB | pUT18 containing drt*opoIB,* $Amp^R$ | Kota et al., 2021 |
| 17 | pUTCD4$A_M$ | pUT18 carrying Middle domain of CDS (333 bp to 708 bp) of *drdivIVA*, ~3.4kb, $Amp^R$ | Chaudhary et al., 2021 |
| 18 | pUTD4$A_N$ | pUT18 carrying N-terminal domain of *drdivIVA*, ~3.5kb, $Amp^R$ | Chaudhary et al., 2021 |
| 19 | pVHZGFP | pVHSM containing *drftsZ-gfp*, ~11.7kb, $Spec^R$ | Modi et al., 2014 |
| 20 | pNKFKUD | pNOKOUT containing upstream (~1.2 kb) and downstream (~0.7 kb) sequences of dr*ftsK*, ~5.9 kb, $Amp^R$/ $Kan^R$ | This study |
| 21 | pNKFK$_{MC}$UD | pNOKOUT containing upstream sequence (~1.0 kb) of middle region of *drftsk* (884-3627 base) and | This study |

| | | | |
|---|---|---|---|
| | | downstream (~0.8 kb) sequence of *drftsk*, ~5.8 kb, Amp$^{R}$/ Kan$^{R}$ | |
| 22 | pNKFK$_{N}$UD | pNOKOUT containing upstream sequence (~1.0 kb) of N-terminal of *drftsk*(1-489 base) and downstream sequence of N-terminal of *drftsk* (~0.8 kb) , ~5.8 kb, Amp$^{R}$/ Kan$^{R}$ | This study |
| 23 | pDsRedFKγ | pDsRed carrying *drftskγ,* ~3.7kb, Amp$^{R}$ | This study |
| 24 | pNKFKγRD | pNOKOUT containing *drftskγ-rfp* (~1.2 kb) and downstream (~1.2 kb) sequences of *ftsK*, ~6.4kb, Amp$^{R}$/ Kan$^{R}$ | This study |

Table S2: List of primers used in this study.

| **S.No** | **Primer Name** | **Oligonucleotide Sequences** | **Purpose** |
|---|---|---|---|
| 1 | DsRedFkGFw | CGGGATCCGGCAGCTGACATCCTGACCAG C | pDsRedFKγ construction |
| 2 | DsRedFkGRw | GGGGTACCGCGGCCGCTTTGCCGAAATAC TC | |
| 3 | FKGUpFw | GGCGGGCCCGACATCCTGACCAGC | pNKFKγRD construction |
| 4 | RFPUpRw | CGGAATTCCTACAGGAACAGGTG | |
| 5 | FKGDnFw | CGGGATCCGGCGACTGTGGCCGC | for *ftsk-rfp* knock-in mutant |
| 6 | FkGDnRw | GCTCTAGAACGTGGAAGCGCAGC | |
| 7 | FKMCUpFw | CGGGATCCTCCACCACGGCCTTG | pNKFK$_{MC}$U D construction for Δ*ftsKMC* |
| 8 | FKMCUpRw | GCTCTAGACAGTGGATGCGGAGGTAC | |
| 9 | FKDnFw | GGCGGGCCCACGTGGAAGCGCAGCAGCG AG | pNKFK$_{MC}$U D and |
| 10 | FKDnRw | CGGAATTCCGCAAACAAACAGAATTACTG | pNKFKUD construction |
| 11 | FKUpFw | CGGGATCCGT GGTGACCTGTTTTGCAGG | |

| | | | |
|---|---|---|---|
| 12 | FKUpRw | GCTCTAGA GTTAACGAGAGTAGCCGCGAC | pNKFKUD construction for Δ*ftsK* |
| 13 | FKNUpFw | GGC GGGCCC TGCTCTCAGCTGCCG | pNKFK$_N$UD construction for Δ*ftsKN* |
| 14 | FKNUpRw | CGGAATTCGTGCATGGGCTGATC | |
| 15 | FKNDnFw | CGGGATCCCGGTTTAGTTTATAC | |
| 16 | FkNDnRw | GCTCTAGACATGGTGGGCCTGAACGT | |
| 17 | FKFw | CCCATATGATGTCGGCAAACTCG | pETFtsK construction |
| 18 | 0400Fw | CCCATATGGTGCGGCGCCTGCAACGC | pET0400 construction |
| 19 | DFKγFw | CCCATATGGACATCCTGACCAGC | pETDFKγ construction |
| 20 | FKRw | CGGGATCCCTATTTGCCGAAATAC | pETFtsK, pETDFKγ and pET0400 construction |
| 21 | EFKγFw | CCCATATGAGCGAAGGTGGTGCG | pETEFKγ construction |
| 22 | EFKγRw | CGGGATCCTTAGTCAAACGGCGG | |
| 23 | EXCFw | CCCATATGATGACCGATTTACAC | pETEXC construction |
| 24 | EXCRw | CGGGATCCTTATTTCCCCCGTTT | |
| 25 | EXDFw | CCCATATGGTGAAACAGGATCTG | pETEXD construction |
| 26 | EXDRw | CGGGATCCTCACGCCCGCGGGTGA | |
| 27 | AF | ATGTCGGCAAACTCG | Diagnostic PCR |
| 28 | AR | CAGCCGCCGGGTCAC | |
| 29 | CF | CTGCCGAGCGAGCAGCTC | |
| 30 | CR | CACGCTGGGCCCGCTG | |
| 31 | NF | GCACGGTGGCCGAGTGG | |
| 32 | NR | GTCAGCGTAATGCTCTG | |
| 33 | MR | GTGGTCGCCTGCTCGCCT | |
| 34 | UF | AGGGTCAGGAGCACGCCC | |
| 35 | DR | GCCAAATTGCAACACTTGTACCTT | |
